# Disruption of the MICOS complex leads to an aberrant cristae structure and an unexpected, pronounced lifespan extension in *Podospora anserina*

**DOI:** 10.1101/2022.03.21.485166

**Authors:** Verena Warnsmann, Lisa-Marie Marschall, Anja C. Meeßen, Maike Wolters, Lea Schürmanns, Marion Basoglu, Stefan Eimer, Heinz D. Osiewacz

## Abstract

Mitochondria are dynamic eukaryotic organelles involved in a variety of essential cellular processes including the generation of adenosine triphosphate (ATP) and reactive oxygen species (ROS) as well as in the control of apoptosis and autophagy. Impairments of mitochondrial functions lead to aging and disease. Previous works with the ascomycete *Podospora anserina* demonstrated that mitochondrial morphotype as well as mitochondrial ultrastructure change during aging. The latter goes along with an age-dependent reorganization of the inner mitochondrial membrane leading to a change from lamellar cristae to vesicular structures. Especially from studies with yeast it is known that beside the F_1_F_o_-ATP-synthase and the phospholipid cardiolipin also the ‘mitochondrial contact site and cristae organizing system’ (MICOS) complex, existing of the Mic60- and Mic10-subcomplex, is essential for proper cristae formation. In the present study, we aimed to understand the mechanistic basis of age-related changes of the mitochondrial ultrastructure. We observed that MICOS subunits are co-regulated at the posttranscriptional level. This regulation partially depends on the mitochondrial iAAA-protease PaIAP. Most surprisingly, we made the counterintuitive observation that, despite the loss of lamellar cristae and of mitochondrial impairments, the ablation of MICOS subunits (except of PaMIC12) leads to a pronounced lifespan extension. Moreover, simultaneous ablation of subunits of both MICOS subcomplexes synergistically increases lifespan, providing formal genetic evidence that both subcomplexes affect lifespan by different and at least partially independent pathways. At the molecular level we found that ablation of Mic10-subcomplex components leads to a mitohormesis-induced lifespan extension, while lifespan extension of Mic60-subcomplex mutants seems to be controlled by another pathway. Overall, our data demonstrate that both MICOS subcomplexes have different functions and play distinct roles in the aging process of *P. anserina*.

## 1 INTRODUCTION

Mitochondria are double-membrane bound, ubiquitous eukaryotic organelles with a variety of essential functions. They play a central role in organismal aging and human diseases (Breitenbach et al., 2012; Kauppila et al., 2017; Osiewacz, 2010; Sun et al., 2016; Tatsuta & Langer, 2008). The inner mitochondrial membrane forms invaginations, the so-called cristae. At their basis, cristae junctions connect the cristae membrane with the inner boundary membrane, the part of the inner membrane which is closely apposed to the outer membrane (Frey & Mannella, 2000; Klecker & Westermann, 2021; Zick et al., 2009). This characteristic architecture is crucial for mitochondrial function (Mannella, 2006). Cristae morphology is highly variable and was found to change during aging in various organisms and in numerous human disorders (Colina-Tenorio et al., 2020). For instance, in the filamentous fungus *Podospora anserina*, a well-established aging model with a strong mitochondrial etiology of aging, functional mitochondria from young individuals contain lamellar cristae whereas mitochondria from older cultures lose this typical cristae structure and the inner membrane forms vesicles (Daum et al., 2013). The formation and maintenance of cristae are mediated by the interplay of two major protein complexes and the mitochondrial phospholipid cardiolipin (Harner et al., 2016; Klecker & Westermann, 2021; Martensson et al., 2017; Rabl et al., 2009). The first complex is the dimeric F_1_F_o_-ATP-synthase, which is required for the positive curvature at the cristae tips (Bornhövd et al., 2006; Davies et al., 2012; Strauss et al., 2008). Loss of F_1_F_o_-ATP-synthase dimers leads to aberrant cristae structures (Arnold et al., 1998; Habersetzer et al., 2013; Paumard et al., 2002; Rampello et al., 2018). For instance, in *P. anserina* the loss of F_1_F_o_-ATP-synthase dimers causes the loss of lamellar cristae and a mitophagy-dependent reduction of lifespan (Rampello et al., 2018; Warnsmann et al., 2021).

The second protein complex is the evolutionary conserved MICOS complex (‘mitochondrial contact site and cristae organizing system’). This complex is responsible for negative membrane curvature at the base of cristae, for cristae junction formation as well as connections of the inner and outer mitochondrial membrane (Harner et al., 2011; Hoppins et al., 2011; van der Laan et al., 2012; von der Malsburg et al., 2011). The MICOS complex is composed of several subunits. In the budding yeast, MICOS contains six subunits, which are organized in two subcomplexes, a Mic60- and a Mic10-subcomplex (Figure 1A). The Mic60-subcomplex consists of the core-subunit MIC60 and the regulatory subunit MIC19, and connects the inner and outer mitochondrial membranes. The Mic10-subcomplex of yeast is composed of the core-subunit MIC10 und the regulatory subunits MIC26, MIC27 and MIC12. Oligomerization of MIC10 monomers initiates membrane curvature at cristae junctions. In the human MICOS complex the Mic60-subcomplex contains MIC25 as an additional subunit. Furthermore, instead of MIC12, MIC13/QIL1 is part of the human Mic10-subcomplex. The loss of individual MICOS subunits causes a disruption of cristae junctions combined with aberrant cristae structure (Khosravi & Harner, 2020; Rampelt et al., 2017; Wollweber et al., 2017).

**Figure 1:**
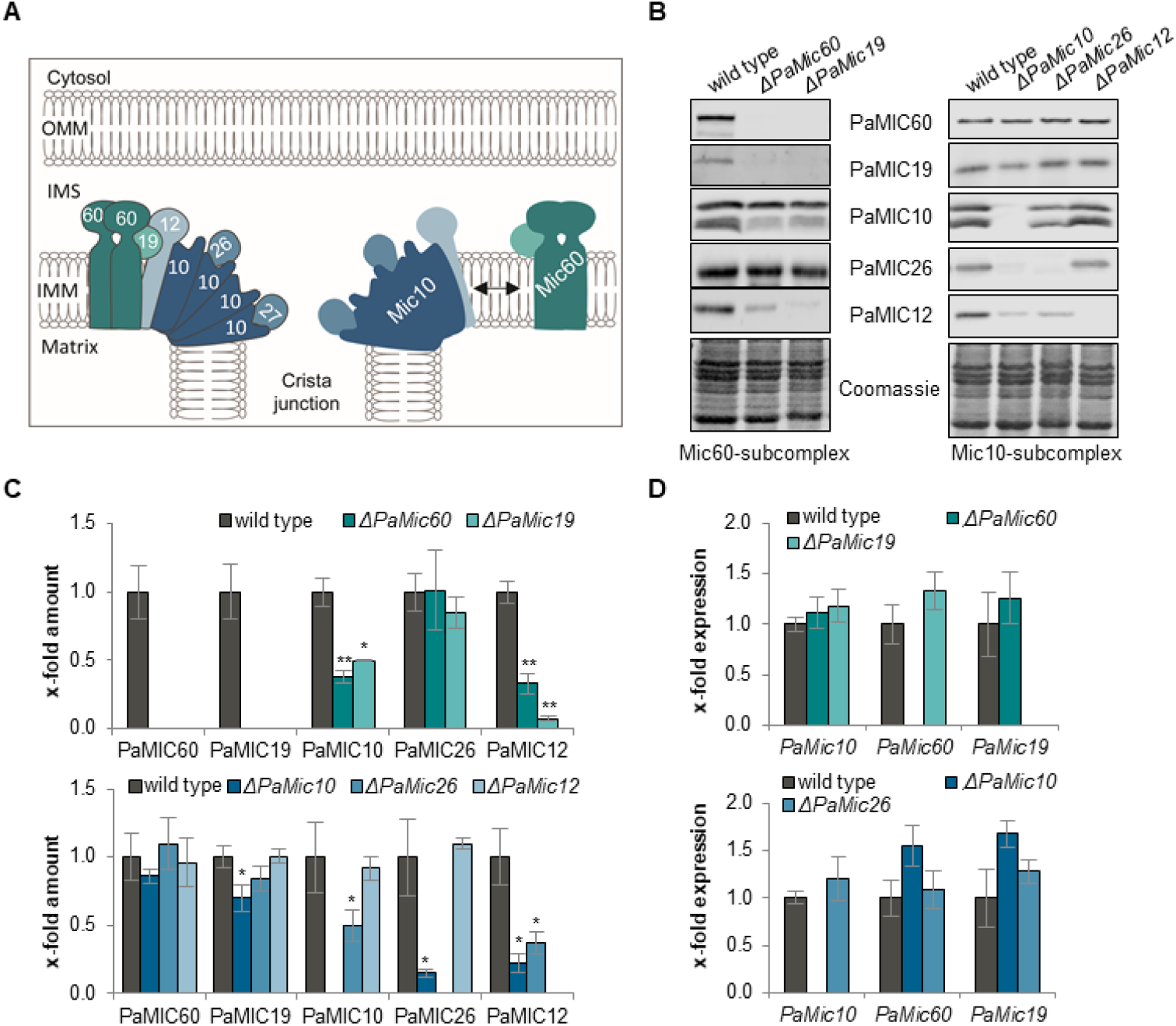
MICOS subunits affect each other. (A) Scheme depicting the MICOS subcomplexes and the individual subunits in yeast. The Mic60-subcomplex consists of the subunits MIC60 and MIC19. MIC10, MIC27, MIC26 and MIC12 form the Mic10-subcomplex. Together both subcomplexes form the MICOS complex. OMM: outer mitochondrial membrane; IMS: intermembrane space; IMM: inner mitochondrial membrane. (B) Western blot analysis of mitochondrial protein extracts of *P. anserina* wild type, *ΔPaMic60, ΔPaMic19, ΔPaMic10, ΔPaMic26* and *ΔPaMic12* using antibodies against the indicated proteins. (C) Quantification of MICOS protein abundance normalized to the Coomassie-stained gel and related to the wild type (set to 1). Data are mean abundance ± SD (each 3 biological replicates). (D) Relative transcript levels of *PaMic10, PaMic60* and *PaMic19* in wild type, *ΔPaMic60, ΔPaMic19* and wild type, *ΔPaMic10* and *ΔPaMic26* (each 3 biological replicates). Transcript levels of wild-type cultures were set to 1. Data are mean expression ± SD. * Significant differences to wild type; \**p* < 0.05; \*\**p* < 0.01.

Both cristae-forming protein complexes, F_1_F_o_-ATP-synthase dimers and the MICOS complex, are stabilized by the mitochondrial phospholipid cardiolipin, which promotes cristae formation (Acehan et al., 2011; Friedman et al., 2015; Hoppins et al., 2011; Martensson et al., 2017). Moreover, due to the cone-shaped structure, cardiolipin itself is involved in cristae curvature formation (Beltran-Heredia et al., 2019; Khalifat et al., 2008). In yeast, plants and flies mutants lacking cardiolipin synthase (CRD1) or the cardiolipin remodeling enzyme tafazzin (TAZ1) cristae structure is abnormal (Acehan et al., 2011; Acehan et al., 2007; Baile et al., 2014; Claypool et al., 2008; Pan et al., 2014; Pineau et al., 2013; Xu et al., 2006). Similar to impairments in F_1_F_o_-ATP-synthase dimerization, ablation of CRD1 leads to lifespan reduction in *Drosophila melanogaster* and *P. anserina* (Acehan et al., 2011; Löser et al., 2021).

Although the structure of MICOS and its importance for the mitochondrial ultrastructure are well studied in yeast and human, the functional impact of MICOS impairments on the aging process remains unclear. For *P. anserina*, a previously published model postulates that the MICOS complex has to dissociate during aging to allow the transition from tubular to vesicular inner membrane organization (Daum et al., 2013). In the current study, we set out to experimentally address this process in more detail. First, we identified homologs to the yeast MICOS subunits in *P. anserina* and analyzed their regulation and impact on organismic aging. We demonstrate that the steady-state levels of identified MICOS subunits in *P. anserina* are co-regulated, and this regulation is partially dependent on mitochondrial iAAA-protease PaIAP. Surprisingly, we made the counterintuitive observation that, despite the defect of cristae junction formation, the ablation of MICOS subunits leads to a pronounced lifespan extension. So far, the MICOS complex was thought to act as a unit. However, in contrast here we demonstrate that the MICOS complex rather acts as two subcomplexes which both regulate lifespan through different pathways. In Mic10-subcomplex mutants the lifespan extension results from the induction of mitohormesis, which does not appear to play a major role in lifespan extension of Mic60-subcomplex mutants.

## 2 MATERIALS AND METHODS

### 2.1 *P. anserina* strains and cultivation

In this study, the *P. anserina* wild-type strain ‘s’ (Rizet, 1953), *PaSod3*^*H26L*^::*Gfp* (Knuppertz et al., 2017), *ΔPaIap* (Weil et al., 2011) as well as the newly generated mutants *ΔPaMic60, ΔPaMic19, ΔPaMic10, ΔPaMic26, ΔPaMic12, ΔPaMic60/ΔPaIap, ΔPaMic10/ΔPaIap, ΔPaMic26/ΔPaIap ΔPaMic60/PaSod3*^*H26L*^::*Gfp, ΔPaMic19/PaSod3*^*H26L*^::*Gfp, ΔPaMic10/PaSod3*^*H26L*^::*Gfp, ΔPaMic26/PaSod3*^*H26L*^::*Gfp* and *ΔPaMic12/PaSod3*^*H26L*^::*Gfp* were used. Strains were grown on standard cornmeal agar (BMM) at 27 °C under constant light (Osiewacz et al., 2013). For spore germination BMM with 60 mM ammonium acetate (Merck, Darmstadt, Germany; 1116.1000) was used. Spores were incubated at 27 °C in the dark for 2 days. All strains used in this study were derived from monokaryotic ascospores (Osiewacz et al., 2013).

### 2.2 Cloning procedures and generation of *P. anserina* mutants

All transgenic strains are in the genetic background of wild-type strain ‘s’. MICOS deletions strains were generated according to a method developed by El-Khoury and colleagues using the plasmid pKO7 and the strain *ΔPaKu70* (El-Khoury et al., 2008; Kunstmann & Osiewacz, 2009). In short, approximately 1 kbp long fragments corresponding to the 5’ and 3’ flanking regions of the respective gene of interest were amplified by PCR using sequence specific oligonucleotides (Table S1) with appropriate restriction site overhangs and cloned into the plasmid pKO7 to flank a hygromycin B resistance gene. For deletion of *PaMic26, PaMic12* and *PaMic19* a cloning strategy with the restriction sites ClaI and HindIII (5’-region) and PstI and BamHI (3’-region) was used. In case of *PaMic10* and *PaMic60* different cloning strategies were used. The 5’-regions were cloned with the restrictions sites BamHI and PstI (*PaMic60*) or KpnI and HindIII (*PaMic10*). For cloning of 3’-regions the restriction enzymes EcoRV and HindIII (*PaMic60*) or PstI and BamHI (*PaMic10*) were used. The resulting gene specific deletion vectors were used to transform *P. anserina* spheroplasts of the phleomycin resistant *ΔPaKu70* strain. Transformants were selected by their hygromycin B resistance and crossed with the wild type to reintroduce the *PaKu70* gene. Progenies of these crosses were selected by their hygromycin B resistance and phleomycin sensitivity and further verified by Southern blot analysis.

For complementation analysis MICOS genes with promotor, terminator and appropriate restriction site overhangs were amplified with specific oligonucleotides (Table S2) and cloned into the plasmid pKO6 containing a phleomycin resistance cassette (Knuppertz et al., 2014). After transformation of the resulting plasmid in spheroplasts of the corresponding deletion strain, selection of transformants was performed by growth on phleomycin containing medium.

### 2.3 Generation of *P. anserina* double mutants

For double mutant generation the single mutant strains were crossed with each other. Subsequently, strains were selected from the progeny containing both mutations.

### 2.4 Southern blot analysis

DNA isolation was performed according to a well-established protocol from Lecellier and Silar (Lecellier & Silar, 1994). From each sample 500 ng DNA was digested with EcoRV. DNA digestion, gel electrophoresis and Southern blotting was carried out by standard protocols. According to the manufacturer’s protocol for Southern blot hybridization and detection digoxigenin-labeled hybridization probes (DIG DNA Labeling and Detection Kit, Roche Applied Science, 11175033910) were used. The phleomycin resistance gene (*Ble*) specific hybridization probe corresponds to the 1293 bp BamHI-fragment of the plasmid pKO4 (Luce & Osiewacz, 2009). The 736 bp XhoI-fragment of the plasmid pSM4 (Zintel et al., 2010) was used as a specific hybridization probe for the hygromycin resistance gene (*Hyg*). The specific hybridization probes of the MICOS genes were amplified by PCR using gene specific oligonucleotides (Table S3).

### 2.5 Growth rate and lifespan determination

Determination of the lifespan and the growth rate of *P. anserina* cultures derived from monokaryotic ascospores was performed on M2 medium at 27 °C and constant light as previously described (Osiewacz et al., 2013). The lifespan of *P. anserina* is defined as the time period in days (d) until growth stops. The growth rate is defined as the distance of growth (cm) of a culture per time period (d).

### 2.6 Isolation of total protein extracts

Total protein extracts of *P. anserina* strains were isolated as previously described (Warnsmann et al., 2021). Briefly, grown mycelia were homogenized in protein isolation buffer containing 5 mM dithiothreitol (DTT) (Carl Roth, Karlsruhe, Germany, 6908.4) and subsequently sedimented at 9300x *g*. The supernatant was used as total protein extracts and stored at -20 °C.

### 2.7 Isolation of mitochondria

For isolation of mitochondria *P. anserina* strains were grown on cellophane foil covered solid M2 agar for two days at 27 °C and constant light. Grown mycelial pieces were transferred to CM-liquid medium and incubated for additional two days at 27 °C and constant light. The mitochondria of *P. anserina* cultures were isolated according to a published protocol (Gredilla et al., 2006). Afterwards, mitochondria were purified with a discontinuous sucrose gradient (20-36-50%) and ultracentrifugation (100000x *g*) (Osiewacz et al, 2013).

### 2.8 Blue native gel electrophoresis (BN-PAGE)

BN-PAGE to separate native protein complexes according to size was performed as described (Wittig et al., 2006). Briefly, 150 μg of isolated mitochondria were solubilized with digitonin (Sigma-Aldrich, D141) at a detergent/protein ratio of 4 g/g. Solubilized mitochondria were separated on a linear gradient gel (4-13%) overlaid with 3.5% stacking gel. After electrophoresis protein complexes were fixed for 30 min in fixing solution (50% methanol, 10% acetic acid) and subsequently proteins were visualized by 1 h Coomassie staining (0.025% Coomassie blue in 10% acetic acid).

### 2.9 Western blot analysis

Western blot analysis was performed with 50 µg mitochondria or total protein extract as previously described (Warnsmann et al., 2021). The following primary antibodies were used: anti-GFP (mouse, 1:10000 dilution, Sigma-Aldrich, G6795), anti-PaIAP (rabbit, dilution 1:2500, peptide: [H]-KAENQKARFSDVHGC-[OH], Sigma-Genosys, Haverhill, United Kingdom), anti-PaMIC10 (rabbit, dilution 1:10000, peptide: [H]-AYEECNSSLKQAAKEIRKQA-[OH], Davids Biotechnologie GmbH, Regensburg, Germany), anti-PaMIC60 (rabbit, dilution 1:10000, peptide: [H]-KAKKETAALPKVEAKDAALEKK-[OH], Davids Biotechnologie GmbH, Regensburg, Germany), anti-PaMIC19 (rabbit, dilution 1:2000, peptide: [H]-REVEAFKEEVRRVEKGWVEK-[OH], Davids Biotechnologie GmbH, Regensburg, Germany), anti-PaMIC26 (rabbit, dilution 1:10000, peptide: [H]-PRKPIYDDDLDDPLPTSK-[OH], Davids Biotechnologie GmbH, Regensburg, Germany), anti-PaMIC12 (rabbit, dilution 1:10000, peptide: [H]-ERALAERFDQAKRERRLERS-[OH], Davids Biotechnologie GmbH, Regensburg, Germany).

As secondary antibodies IRDye^®^ 680RD anti-rabbit (goat, 1:15000 dilution, LIC-OR Biosciences, Bad Homburg, Germany, 926-68071), IRDye^®^ 800CW anti-rabbit (goat, 1:15000 dilution, LIC-OR Biosciences, Bad Homburg, Germany, 926-32211) and IRDye® 680RD anti-mouse (goat, dilution 1:15000, LI-COR Biosciences, Bad Homburg, Germany, 926-68070) were used. For detection the Odyssey^®^ Fc imaging system (LIC-OR Biosciences, Bad Homburg, Germany,) was used and densitometric quantification was performed with the manufacturer’s software image studio (Version 5.2).

### 2.10 Quantitative real-time PCR (qRT-PCR)

RNA isolation and cDNA synthesis was performed as previously described (Heinz et al., 2021). For each strain three biological replicates were analyzed three times each (technical replicates). Transcript levels of *PaMic10, PaMic19* and *PaMic60* were determined and normalized to transcript level of *PaPorin*. The used oligonucleotides are listed in Table S4. For all genes, PCR efficiency was determined as described in (Pfaffl, 2001).

### 2.11 Superoxide and hydrogen peroxide release measurements

For qualitative determination of superoxide anion release from mycelia a histochemical nitro blue tetrazolium (NBT, Sigma-Aldrich, N6876) staining was performed as previously described (Warnsmann et al., 2018a). Analogously, for hydrogen peroxide release a histochemical diaminobenzidine (DAB, Sigma-Aldrich, D-8001) staining was performed as previously described (Warnsmann et al., 2018a)

### 2.12 Fluorescence microscopy

*P. anserina* strains were cultivated on glass slides with a central depression containing 130 µl M2 medium for 1 day under standard conditions. Fluorescence microscopic analyses were performed with the confocal spinning disc microscope Zeiss Cell Observer SD and a 63x/1.4 oil objective lens (Carl Zeiss Microscopy, Jena, Germany) using a 488 nm laser line. Image processing was performed with the corresponding software ZEN 2.5 (blue edition).

### 2.13 Transmission electron microscopy

For transmission electron microscopy, *P. anserina* cultures were grown on M2 agar and small agar blocks were cut out at the 6-day growths front and fixed while shaking for 1 h at room temperature (RT) in 0.1 M cacodylate buffer at pH 7.2 containing 0.25 M sucrose and 4% glutaraldehyde. The fixed samples were then washed two times in 0.1 M cacodylate buffer pH 7.2 containing 10% sucrose at RT. Afterwards the samples were stained with 2% OsO4 for 3 h at RT and washed twice with wash buffer. Prior to embedding samples were dehydrated by using increasing concentrations of ethanol buffer from 50%, 70%, 90% and twice 100%, followed by two washes with propylene oxide. Subsequently, samples were embedded in Araldite resin and polymerized. 50 nm thin sections were cut using a Reichert Ultracut ultramicrotome and transferred to coated copper slot grids. Grids were post-stained with uranyl acetate and lead citrate and analyzed using a digitalized Zeiss TEM 900 operated at 80 keV and equipped with a Troendle 2K camera.

### 2.14 Statistical Analysis

Significances of differences in lifespan of individual cultures were statistically analyzed with the IBM SPSS statistics 19 software package (IBM, Armonk, NY, USA) by generating the Kaplan-Meier survival estimates. We use three independent statistical tests (Breslow (generalized Wilcoxon), log-rank (Mantel-Cox), and the Tarone-Ware) with a pairwise comparison. All other statistical significances were calculated with the Student’s *t*-test. The respective samples were compared with the appropriate wild-type sample. For statistical significance the minimum threshold was set at p ≤ 0.05. * *p* ≤ 0.05; ** *p* ≤ 0.01; *** *p* ≤ 0.001.

## 3 RESULTS AND DISCUSSION

### 3.1 Identification of MICOS homologs in *P. anserina*

Various studies from yeast to human revealed the structure of MICOS complex and its role for the mitochondrial ultrastructure. Nevertheless, the functional impact of MICOS impairments on the aging process, remains unclear. For *P. anserina*, a previously published model postulates that the MICOS complex has to dissociate during aging to allow the age-dependent reorganization of inner membrane (Daum et al., 2013). In the current study, we set out to experimentally address this process in more detail. Since, previous studies demonstrated that the MICOS complex is evolutionary conserved (Munoz-Gomez et al., 2017), we first performed a protein BLAST search (http://podospora.i2bc.paris-saclay.fr/blast.php) and identified five homologs of the six yeast MICOS subunits (Table 1, Figure S1). We termed them PaMIC60, PaMIC19, PaMIC10, PaMIC26 and PaMIC12 in accordance with the published uniform nomenclature (Pfanner et al., 2014). All identified *P. anserina* MICOS subunits possess the typical domains, which are known from yeast and humans (Table 1). For instance, the core subunit PaMIC60 contains a mitofilin domain that is conserved among organisms (Table 1; Figure S1A). Furthermore, the sequence of PaMIC26 shows the same apolipoprotein-O domain as the corresponding protein in *Saccharomyces cerevisiae* (Table 1). A homolog of the sixth *S. cerevisiae* MICOS subunit MIC27 was not found in *P. anserina*.

**Table 1:** Identification of homologs of the *S. cerevisiae* MICOS complex in *P. anserina*. Putative homologs were identified in a Blast search or based on domain conservation in InterPro database using *S. cerevisiae* as query. UniProt and InterPro accession numbers are provided in brackets. The *P. anserina* MICOS proteins (trivial name) were termed according to the published nomenclature (Pfanner et al., 2014). The numbers correspond to the molecular mass of the proteins.

| MICOS subunit | *S. cerevisiae* | conserved domain | put. homolog in *P. anserina* | trivial name |
| --- | --- | --- | --- | --- |
| MIC60 | YKR016W (P36112) | Mitofilin (IPR019133) | Pa_1_1530 (B2A9R4) | PaMIC60 |
| MIC19 | YFR011C (P43594) | DUF1690 (IPR012471) | Pa_1_19620 (B2AUM5) | PaMIC19 |
| MIC10 | YCL057C-A (Q96VH5) | DUF543 (IPR007512) | Pa_7_7950 (B2AWQ3) | PaMIC10 |
| MIC12 | YBR262C (P38341) | AIM5 (IPR031463) | Pa_2_1415 (B2B416) | PaMIC12 |
| MIC26 | YGR235C (P50087) | Apolipoprotein-O (IPR019166) | Pa_1_10590 (B2AYB9) | PaMIC26 |
| MIC27 | YNL100W (P50945) | no | no | no |

### 3.2 PaIAP-dependent regulation of MICOS steady-state levels

To experimentally address the question whether or not the MICOS complex plays a role in *P. anserina* aging, as it was suggested in an earlier study (Daum et al., 2013), we generated five *P. anserina* strains each lacking one of the individual MICOS subunits. The corresponding genes were replaced through homologous recombination by a hygromycin resistance cassette. The resulting knockout strains were verified by Southern blot analysis (Figure S2A). Previous studies in yeast demonstrated that loss of a single MICOS subunit can affect the stability/steady-state levels of the other MICOS subunits, suggesting a co-regulation of the MICOS complex subunits (Bohnert et al., 2015; Friedman et al., 2015; Harner et al., 2011; von der Malsburg et al., 2011). To test whether or not this kind of co-regulation also occurs in *P. anserina*, we investigated the steady-state levels of all MICOS subunits in the different *P. anserina* MICOS deletion strains. Indeed, in many cases either complete loss or severe reduction of the abundance of MICOS subunits occurs in single deletion mutants (Figure 1B,C). In particular, ablation of one of the two Mic60-subcomplex components, PaMIC60 or PaMIC19, leads to the loss of the other subcomplex component. This is similar to the mammalian system where knockdown of *Mic19* leads to the complete loss of Mic60-subcomplex in cell lines (Darshi et al., 2011; Li et al., 2016). In *P. anserina*, proteins of the Mic10-subcomplex are only partially reduced in deletion mutants of the Mic60-subcomplex. In particular, PaMIC10 and PaMIC12 are reduced in both Mic60-subcomplex mutants, whereas PaMIC26 is not affected. Noticeably, two signals were detected with the PaMIC10 antibody which are absent in the *PaMic10* deletion mutant. The upper band probably corresponds to the unprocessed PaMIC10 with mitochondrial targeting sequence (MTS) at the outside of the mitochondria and the lower band reflects PaMIC10 protein after import and processing. Analysis of the Mic10-subcomplex mutants also revealed strong reduction of Mic10-subcomplex proteins other than those deleted in the mutant and minor effects on the Mic60-subcomplex (Figure 1B,C). For instance, ablation of PaMIC10 led to an almost complete loss of PaMIC26 and PaMIC12 and only slight reductions of the Mic60-subcomplex components PaMIC60 and PaMIC19. Ablation of PaMIC26 reduced the other Mic10-subcomplex components but not those of the Mic60-subcomplex. In contrast, in the *PaMic12* deletion mutant the other MICOS components are not affected (Figure 1B,C). This exception was also described in a yeast *Mic12* deletion strain (Harner et al., 2011). Our data suggest that each subcomplex regulates the stability of the other. The loss of one of the two MICOS core-subunit (PaMIC10 or PaMIC60) has the strongest impact on the whole MICOS complex in *P. anserina*. This is in contrast to results from human cells in which only depletion of MIC60, but not MIC10, results in the absence/reduction of all MICOS subunits (Stephan et al., 2020). In yeast, there are inconsistent data reported. While some studies show that the subcomplexes co-regulate each other, others only describe regulation of components of the genetically modified subcomplex and not from the other subcomplex (Bohnert et al., 2015; Friedman et al., 2015; Harner et al., 2011; von der Malsburg et al., 2011). From our data, we conclude that the five MICOS proteins in *P. anserina* cooperate in an intricate manner and are controlled by a regulatory network.

To analyze at which level the regulation of MICOS proteins takes place, we first investigated the transcript level of selected MICOS genes (Figure 1D). In contrast to protein abundance, transcript levels are not changed in MICOS deletion mutants compared to the wild type. Therefore, regulation must occur at the protein level. From a study with mouse embryonic fibroblasts it is known, that in *Mic19* knockdown cells the loss of MIC60 is rescued by simultaneous ablation of iAAA protease YME1L (Li et al., 2016) suggesting that MICOS components are possible substrates of this protease. To examine this possibility in *P. anserina*, we first analyzed the amount of PaIAP, the *P. anserina* homolog of YME1L, in the MICOS deletion mutants (Figure 2A,B). PaIAP was found to be increased in Mic60-subcomplex mutants but not in Mic10-subcomplex mutants. Thus, the iAAA protease is potentially involved at least in the regulation of Mic60-subcomplex abundance in *P. anserina*. To investigate this possibility in more detail, we generated double mutants which concomitantly lack one MICOS subunit and PaIAP (Figure S2B). Investigations of MICOS steady-state levels in these double deletion mutants strongly indicate a partial PaIAP-dependent regulation of the MICOS subunits (Figure 2C-E). In particular, in *ΔPaMic60* the other Mic60-subcomplex component PaMIC19 is not regulated by PaIAP, while the reduction of the core subunit PaMIC10 of the Mic10-subcomplex is partially PaIAP-dependent (Figure 2C,D). Analysis of Mic10-subcomplex mutants revealed that loss of components from both subcomplexes is at least partially PaIAP-dependent. In *ΔPaMic10* the protein abundance of PaMIC19 and PaMIC26 is increased after simultaneous ablation of PaIAP indicating a regulation by the PaIAP protease (Figure 2C,E).

**Figure 2:**
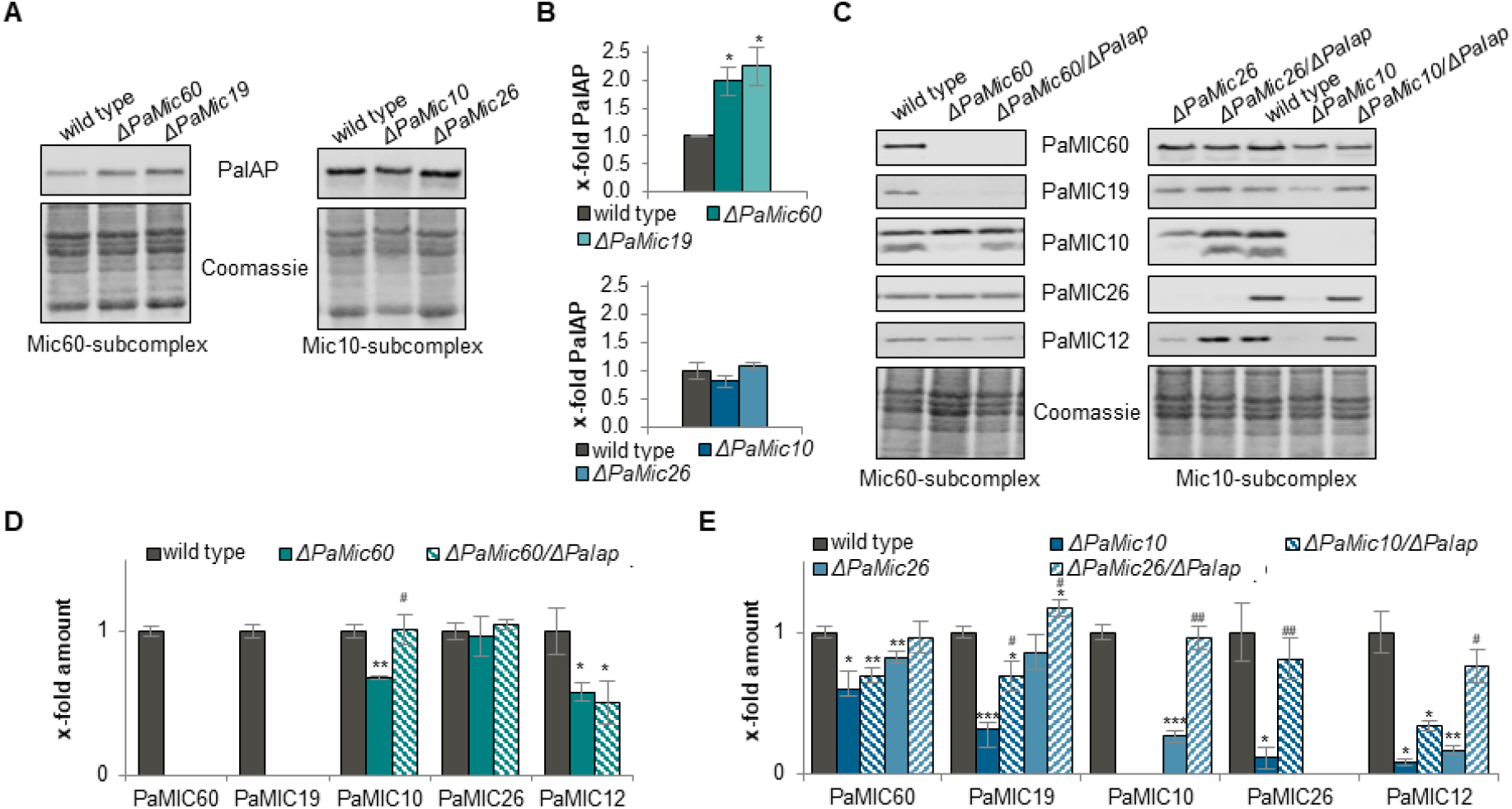
PaIAP-dependent regulation of MICOS steady-state levels. (A) Representative western blot analyses of mitochondrial protein extracts of *P. anserina* wild type, *ΔPaMic60, ΔPaMic19, ΔPaMic10* and *ΔPaMic26* using a PaIAP antibody. (B) Quantification of PaIAP abundance normalized to the Coomassie-stained gel and related to the wild type (set to 1). Data are mean abundance ± SD (each 3 biological replicates). (C) Representative western blot analyses of mitochondrial protein extracts of *P. anserina* wild type, *ΔPaMic60* and *ΔPaMic60/ΔPaIap, ΔPaMic10* and *ΔPaMic10/ΔPaIap, ΔPaMic26* and *ΔPaMic26/ΔPaIap*. MICOS proteins were detected by specific antibodies. (D-E) Quantification of MICOS protein levels. MICOS protein levels were normalized to the Coomassie-stained gel and related to the wild type (set to 1). Data represent mean ± SD (n = 3). * Significant differences to wild type; ^#^ significant differences to the corresponding single MICOS deletion mutant. *^/#^*p* < 0.05; **^/##^*p* < 0.01; ***^/###^*p* < 0.001.

Likewise, the abundance of PaMIC10, PaMIC12 and PaMIC19 in *ΔPaMic26* strains is PaIAP-dependent (Figure 2C,E). Our findings are in agreement with potential substrates of the iAAA protease from different organisms which were identified by substrate trapping assays. For instance, in *Arabidopsis thaliana* and *D. melanogaster* MIC60 was found as a potential substrate of the iAAA protease YME1 (Opalinska et al., 2017; Yoon et al., 2019). Moreover, in yeast, MIC27, MIC10, MIC12 and MIC60 were identified as potential YME1 substrates (Schreiner et al., 2012).

Taken together, our findings extend the current knowledge about the YME1/IAP-dependent regulation of MICOS complex. Hitherto only a YME1-dependent regulation of MIC60 was known and now we demonstrated that the regulation of all MICOS subunits is partially mediated by the iAAA protease PaIAP. Such a proteolytic post-translational regulation allows the organism to very quickly respond to environmental changes, like e.g. different nutrient supply.

### 3.3 Mitochondrial morphology and function are impaired by ablation of MICOS subunits

Next, we set out to investigate whether impairments of MICOS have an impact on whole mitochondria. Since MICOS in yeast and humans is essential to maintain mitochondrial ultrastructure (Harner et al., 2011; Stephan et al., 2020; von der Malsburg et al., 2011), we compared mitochondrial cristae organization of MICOS deletion strains and wild-type strains by transmission electron microscopy (TEM) (Figure 3A). Ablation of MICOS in *P. anserina* leads to characteristic alterations of the inner mitochondrial membrane morphology with stacks of cristae membranes and loss of crista junctions. As expected from earlier studies (Brust et al., 2010; Rampello et al., 2018), the wild type contains mostly filamentous mitochondria with short cristae, which are connected to the inner boundary membrane in 95% of the mitochondria (Figure 3A,B). In contrast, in *ΔPaMic60, ΔPaMic19, ΔPaMic10* and *ΔPaMic26* mutants mitochondria are highly fragmented with cristae, which are not connected to the inner boundary membrane in 91%, 79%, 93% and 89% of the mitochondria, respectively (Figure 3A,B). These floating cristae often form membrane stacks parallel to the longest mitochondrial axis. The only exception is the *ΔPaMic12* mutant, which mostly shows filamentous mitochondria with only a small fraction of about 16% of abnormal mitochondria. Similar, rather wild-type-like cristae structure was also described for yeast *Mic12* deletion strains (Harner et al., 2011).

**Figure 3:**
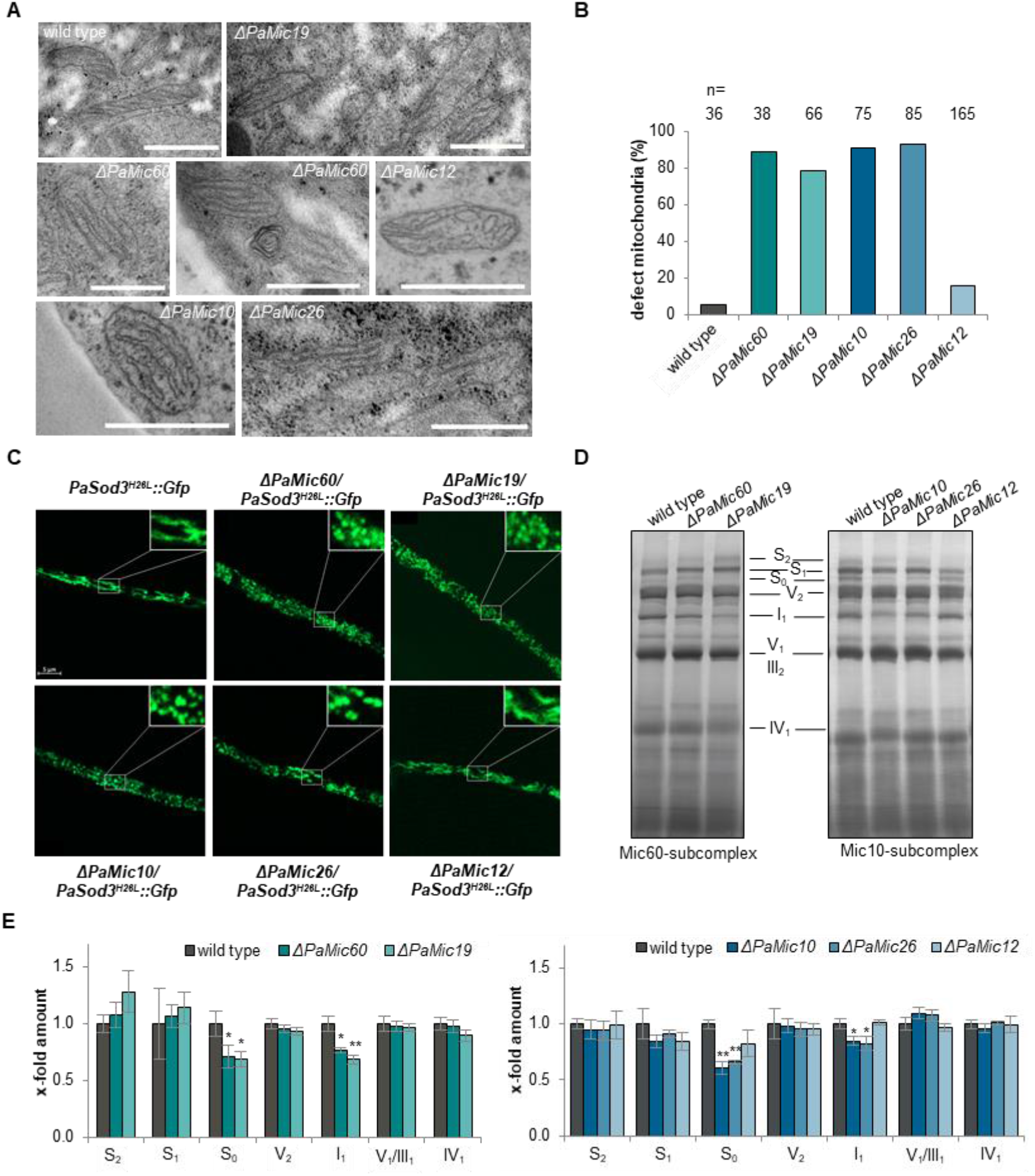
Ablation of MICOS subunits leads to mitochondrial impairments. (A) Representative TEM images of mitochondria from 50 nm thin sections of plastic embedded 6-day-old *P. anserina* mycelia of wild-type and MICOS deletion strains. Scale bar: 250 nm (B) Quantification of the number of aberrant mitochondria with floating cristae in *P. anserina* wild type and MICOS mutants. n = number of mitochondria analyzed. (C) Fluorescence microscopic analysis of mitochondria from 6-day-old *P. anserina* wild-type and *ΔPaMic60, ΔPaMic19, ΔPaMic10, ΔPaMic26* and *ΔPaMic12* strains with the mitochondrial reporter PaSOD3^H26L^::GFP. (D) Representative BN-PAGE analysis and (E) quantification of isolated mitochondria from *P. anserina* wild-type and *ΔPaMic60, ΔPaMic19, ΔPaMic10, ΔPaMic26* and *ΔPaMic12* cultures (3 biological replicates for each culture). The supercomplexes (S_0-2_; I_1_III_2_IV_0-2_), dimeric complexes III and V (III_2_ and V_2_) as well as monomeric complexes I_1_, IV_1_, and V_1_ were visualized by Coomassie staining. Data represent mean ± SD. \**p* < 0.05; \*\**p* < 0.01.

In contrast, for ablation of the human MIC12 homolog MIC13 severe changes of the cristae such as in the other MICOS mutants were described (Stephan et al., 2020). The ultrastructural changes observed in *P. anserina* MICOS mutants (except for PaMIC12) are in concordance with findings from yeast and humans, which also describe characteristic cristae stacks (Friedman et al., 2015; Harner et al., 2011; Stephan et al., 2020; von der Malsburg et al., 2011). Nevertheless, these ultrastructural changes in *P. anserina* MICOS mutants were clearly different from the known alterations of the ultrastructure in F_1_F_o_-ATP-synthase mutants or in senescent *P. anserina* wild-type strains. In the senescent *P. anserina* wild type the cristae formed a reticulate network resembling fishing nets (Brust et al., 2010; Daum et al., 2013). A similar reticular or vesicular morphology of the cristae was also shown in *P. anserina* F_1_F_o_-ATP-synthase dimerization mutants (Rampello et al., 2018). These differences in cristae structures of MICOS and F_1_F_o_-ATP-synthase dimerization mutants were also described in yeast and explained by the different roles of these two complexes in cristae formation (Rabl et al., 2009). Accordingly, the MICOS complex is essential for the formation of cristae junctions and the F_1_F_o_-ATP-synthase dimers are necessary for the formation of cristae tips. Hence, F_1_F_o_-ATP-synthase dimerization mutants cannot form cristae tips, leading to bridge-like cristae that extend across whole sections of mitochondria, ending on both sides with fused cristae junctions. Inner membranes forming vesicles in the old *P. anserina* wild type appear to result from the same mechanism as those resulting from the loss of F_1_F_o_-ATP-synthase dimers in strains lacking dimer assembly factors (Daum et al., 2013; Rampello et al., 2018). In contrast, in yeast MICOS mutants the F_1_F_o_-ATP-synthase dimers help shaping the cristae tips but no cristae junctions can be formed to connect the cristae to the inner boundary membrane leading to floating cristae stacks (Rabl et al., 2009). A similar mechanism most likely also accounts for the floating cristae stacks in *P. anserina* MICOS mutants.

To assess whether these ultrastructural changes are accompanied by altered mitochondrial morphology, we performed fluorescence microscopy analysis. Therefore, we generated MICOS deletion strains expressing the gene coding for a specific mitochondrial marker protein, PaSOD3^H26L^::GFP (Knuppertz et al., 2017) (Figure S2C,D). We found that, in contrast to the wild type, in which mitochondria are mainly filamentous, the MICOS mutants of both subcomplexes exhibit fragmented mitochondria (Figure 3C). In concordance with the data from the ultrastructure analysis, the *PaMic12* deletion strain is an exception and shows wild-type-like filamentous mitochondria. Our findings are in contrast to those from human cells, in which only the Mic60-but not the Mic10-subcomplex is crucial to maintain the mitochondrial network (Stephan et al., 2020). However, in MIC10-depleted yeast cells the mitochondrial network is frequently fragmented (Alkhaja et al., 2012).

Next, we investigated whether or not these morphological and ultrastructural changes in the MICOS mutants affect the composition of the respiratory chain. BN-PAGE analyses revealed that, compared to the wild type, the MICOS mutants of both subcomplexes showed only slight changes in the respiratory chain composition (Figure 3D). The amount of supercomplex S_0_ and complex I are significantly reduced in Mic60-subcomplex mutants (Figure 3D,E). These complexes of the respiratory chain are also reduced in *PaMic10* and *PaMic26* deletion mutants (Figure 3D,F). All other complexes, such as F_1_F_o_-ATP-synthase dimers (V_2_), are not altered compared to wild type. As in our previous analyses, in the *PaMic12* deletion mutant no changes in the composition of respiratory chain are present. This is consistent with the observation that deletion of MICOS subunits in HeLa cells has only a modest impact on the composition and function of the respiratory chain (Stephan et al., 2020).

Taken together, the loss of MICOS leads to an altered ultrastructure and morphology of mitochondria and affects composition of the respiratory chain, suggesting that mitochondrial function is impaired.

### 3.4 Ablation of MICOS subunits leads to unexpected longevity of *P. anserina*

Next, we set out to investigate the consequences of the loss of MICOS for *P. anserina*. Phenotypic comparisons of strains revealed minor differences between the MICOS deletion mutants (except of *ΔPaMic12*) and the wild type (Figure 4A). Deletion mutants of the core subunits PaMIC60 and PaMIC10 show reduced rhythmic growth, exhibit a decreased pigmentation, and an increased formation of aerial hyphae. The deletion mutants of the regulatory subunits PaMIC19 and PaMIC26 show a similar but weaker phenotype. As in yeast (Harner et al., 2011), the growth rates of all MICOS mutants are slightly reduced compared to the wild type (Figure 4B,E).

**Figure 4:**
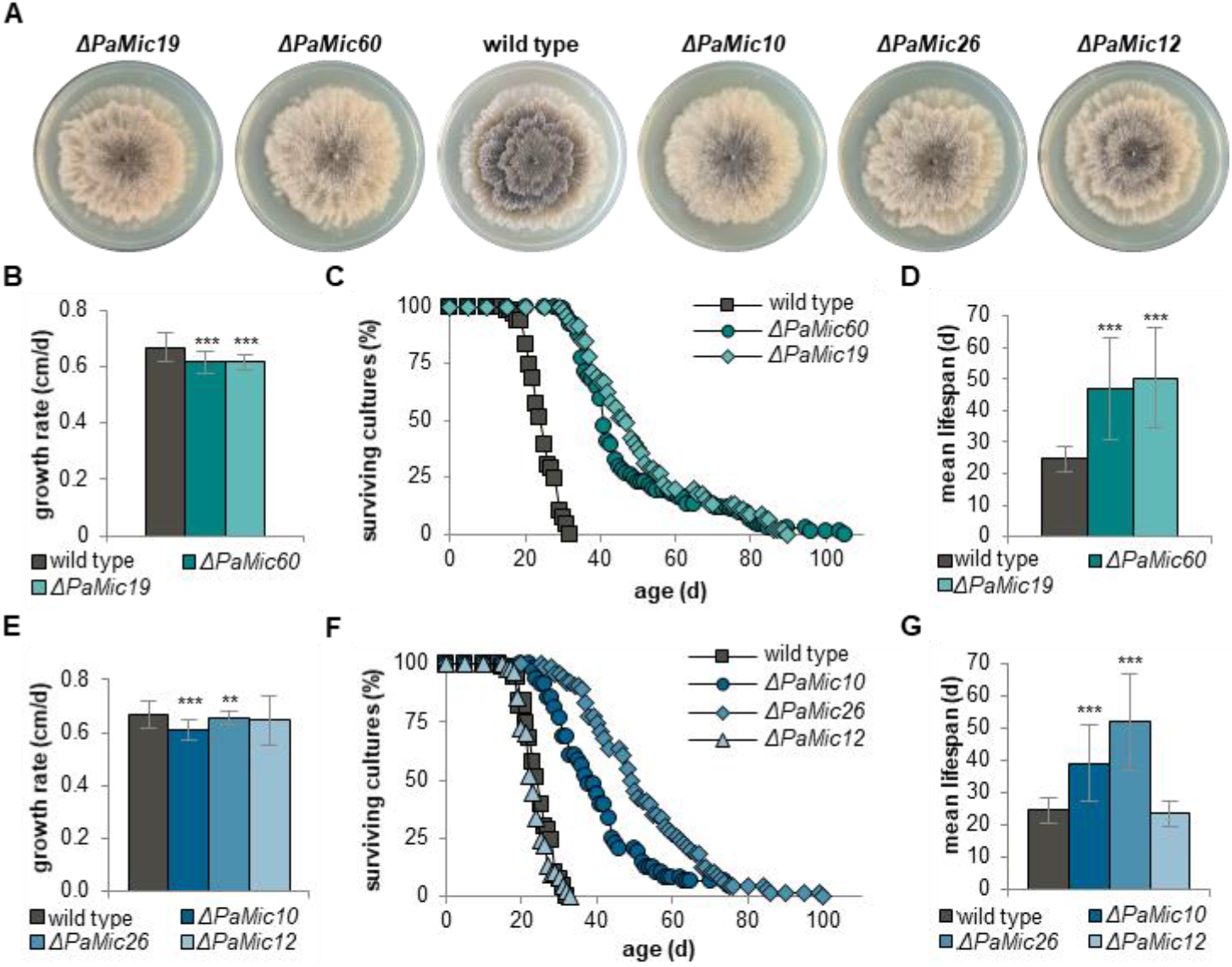
Ablation of MICOS leads to unexpected longevity. (A) Phenotype of 6-day-old *P. anserina* wild-type and *ΔPaMic60 ΔPaMic19, ΔPaMic10, ΔPaMic26* and *ΔPaMic12* strains grown on standard BMM medium. (B) Growth rate and (C) survival curves of *P. anserina* wild type (n = 66); *ΔPaMic60* (n = 74) and *ΔPaMic19* (n = 45) grown on standard M2 medium. (D) Mean lifespan of cultures from (C). (E) Growth rate and (F) survival curves of *P. anserina* wild type (n = 66), *ΔPaMic10* (n = 38), *ΔPaMic26* (n = 66) and *ΔPaMic12* (n = 54) grown on standard M2 medium. (G) Mean lifespan of cultures from (F). Data represent mean ± SD. \*\**p* < 0.01; \*\*\**p* < 0.001.

While the impact of F_1_F_o_-ATP-synthase on mitochondrial ultrastructure and aging of *P. anserina* has been investigated experimentally, such an analysis was missing for MICOS until now. Fragmentation of mitochondrial network and loss of mitochondrial cristae structure are well-known as senescent markers in various organisms, including *P. anserina* (Beregi et al., 1988; Brandt et al., 2017; Brust et al., 2010; Crane et al., 2010; Daum et al., 2013; McQuibban et al., 2006; Scheckhuber et al., 2007; Young et al., 2011; Zhao et al., 2014). Further, we know that fragmentation of mitochondria, due to genetic manipulations or treatment with polyphenols, is accompanied by lifespan reduction in *P. anserina* (Rampello et al., 2018; Warnsmann et al., 2018b). Very recently, a study aiming to develop a new high-throughput replicative lifespan screening with the yeast knockout collection uncovered as a side-effect a decreased replicative lifespan of a *Mic60* deletion strain (Yu et al., 2021). This was a first hint for a possible impact of MICOS on aging. Nevertheless, a detailed analysis of this impact and of the possible basic mechanism is still missing. In our study we analyzed the lifespan of all five *P. anserina* MICOS mutants under standard growth conditions and anticipated to find a lifespan reduction.

Most surprisingly, the ablation of MICOS subunits led to a pronounced lifespan extension (Figure 4C,F; Table S5). The mean lifespan of MICOS mutants is doubled compared to that of wild type (Figure 4D,G). In more detail, ablation of PaMIC60 and PaMIC19 leads to a mean lifespan increase of 88% (47 d vs. 25 d) and 100% (50 d vs, 25 d) and a maximal lifespan extension of 230% (105 d vs. 32 d) and 180% (90 d vs. 32 d), respectively (Figure 4C,F; Table S5). Similarly, ablation of PaMIC10 and PaMIC26 leads to a mean lifespan increase of 56% (39 d vs. 25 d) and 108% (52 d vs. 25) and a maximal lifespan extension of 134% (75 d vs. 32 d) and 213% (100 d vs. 32 d), respectively (Figure 4F; Table S5).

The observed lifespan extension of MICOS mutants is abrogated after the reintroduction of a wild-type copy of the corresponding genes in the MICOS deletion mutants (Figure S3). Again, the *PaMic12* deletion mutant is an exception (Figure 4F). This, together with the observation that in *ΔPaMic12* neither ultrastructure nor the level of other MICOS components are affected, prompts us to speculate about the role of PaMIC12. It might be possible that PaMIC12 is not a central component of the *P. anserina* MICOS. However, the fact that the loss of all other MICOS subunits has an impact on the PaMIC12 steady-state level argues against this possibility (Figure 1B,C). In yeast, MIC12 was shown to function as coupling factor promoting the interaction of the two MICOS subcomplexes (Zerbes et al., 2016). The weak phenotype of *ΔPaMic12* can also be explained by independent roles of the two MICOS subcomplexes, which are not affected upon loss of the coupling factor (Figure 1B,C).

The longevity of MICOS mutants is a counterintuitive finding, since manipulation of the two other cristae-shaping components, F_1_F_o_-ATP-synthase and cardiolipin, has negative effects on organismic aging in *P. anserina* leading to lifespan reduction (Löser et al., 2021; Rampelt et al., 2017). Therefore, we asked how these opposing effects on the lifespan could occur. What might be the physiological reason or the advantages of these differences? Although, future experiments will have to provide the answers to these questions, from our results and the literature, we propose that the reason is the biogenesis of cristae and in the physiological effects of the changes. Various studies demonstrated different ways of cristae formation with different roles of the MICOS complex and the F_1_F_o_-ATP-synthase dimers (Harner et al., 2016; Stephan et al., 2020). Recently, it was postulated that cristae and cristae junctions are highly dynamic undergoing continuously fusion und fission processes (Kondadi et al., 2020). Further, it was shown that re-expression of MICOS proteins in MICOS deletion cell lines could induce secondary cristae junction formation (Stephan et al., 2020). In these studies, the authors assume that cristae stacks in MICOS mutants represent an intermediate structure that also occurs during normal cristae formation in the wild type (Stephan et al., 2020). Hence, we propose that floating cristae in the *P. anserina* MICOS mutants are in some way advantageous compared to the vesicles formed in the F_1_F_o_-ATP-synthase dimerization mutants. Besides the different cristae structure, probably the most important difference is the presence of F_1_F_o_-ATP-synthase dimers in the MICOS mutants which are missing in F_1_F_o_-ATP-synthase dimerization mutants. This difference may lead to different physiological effects in the different mutants.

### 3.5 Increased ROS levels lead to the induction of mitophagy in MICOS mutants

Next, we set out to investigate physiological effects explaining the mechanistic basis of the unexpected lifespan increase of the MICOS mutants. A previous study revealed that mitophagy can act as a pro-survival mechanism in a mitochondrial quality control mutant of *P. anserina* (Knuppertz et al., 2017). To analyze mitophagy in MICOS mutants we performed a biochemical analysis using a vacuolar GFP-cleavage assay (Kanki et al., 2009; Meiling-Wesse et al., 2002). In this assay the vacuolar degradation of the mitochondrial PaSOD3^H26L^::GFP fusion reporter protein is monitored. When PaSOD3^H26L^::GFP labeled mitochondria have entered the vacuole a degradation-resistant GFP-part (‘free GFP’), which was liberated from the fusion protein through proteolytic digestion, can be detected in western blots indicating ongoing mitophagy (Knuppertz et al., 2017). Western blot analysis revealed a nearly 1.5-fold increase in the amount of ‘free GFP’ in the Mic60-subcomplex mutants *ΔPaMic60* and *ΔPaMic19* (Figure 5A,B).

**Figure 5:**
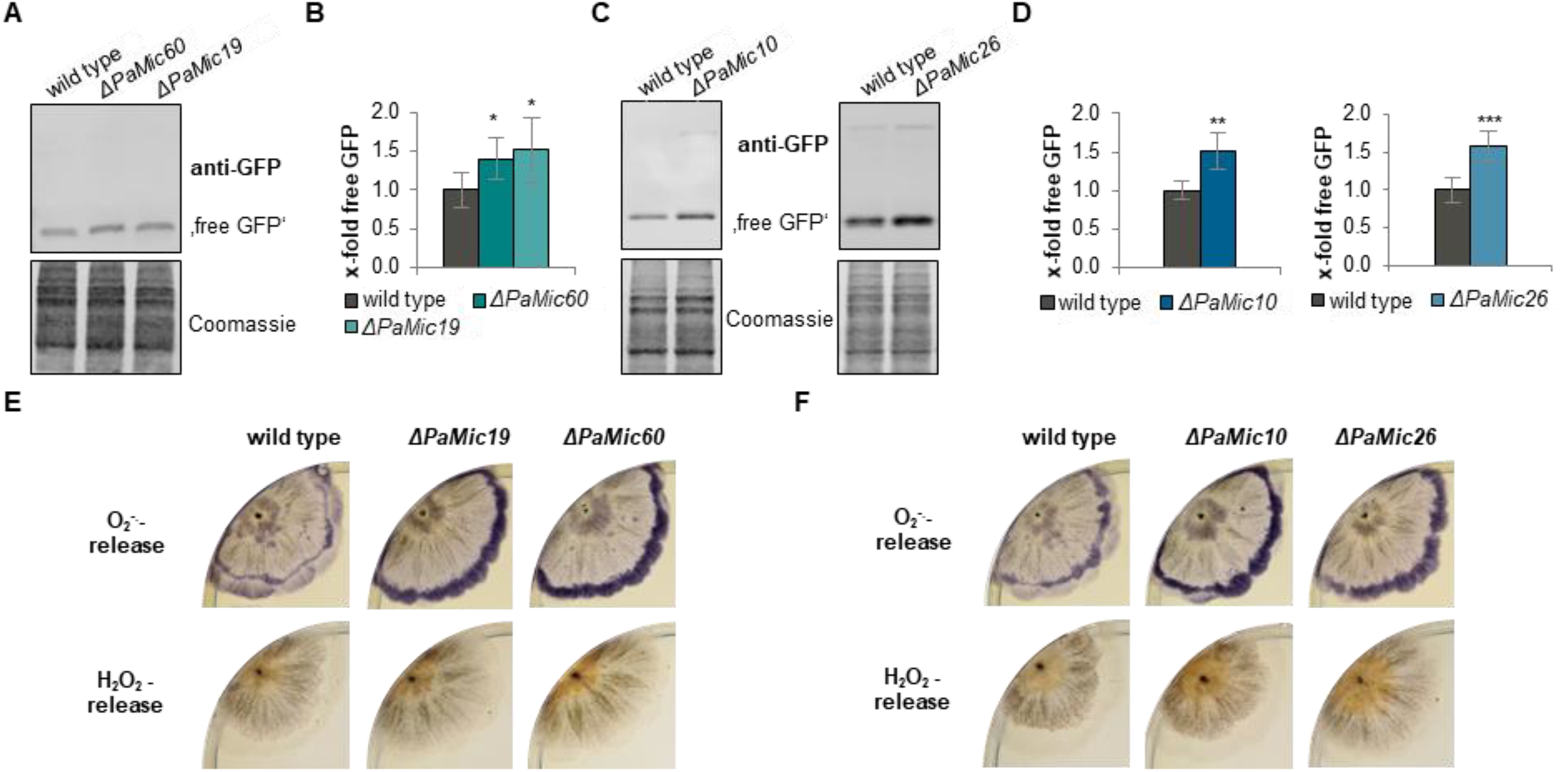
Disruption of MICOS leads to mitophagy induction and oxidative stress. (A) Monitoring mitophagy by western blot analysis of total protein extracts from *PaSod3*^*H26L*^::*Gfp* (here wild type), *ΔPaMic60/PaSod3*^*H26L*^::*Gfp* (here *ΔPaMic60*) and *ΔPaMic19/PaSod3*^*H26L*^::*Gfp* (here *ΔPaMic19*) cultures with a GFP antibody (each 6 biological replicates). (B) Quantification of ‘free GFP’ protein level from (A) normalized to the Coomassie stained gel. Protein level in *PaSod3*^*H26L*^::*Gfp* cultures was set to 1. Data represent mean ± SD. (C) Monitoring mitophagy by western blot analysis of total protein extracts from *PaSod3*^*H26L*^::*Gfp* (here wild type), *ΔPaMic10/PaSod3*^*H26L*^::*Gfp* (here *ΔPaMic10*), and *ΔPaMic26/PaSod3*^*H26L*^::*Gfp* (here *ΔPaMic26*) cultures with a GFP antibody (each 5-7 biological replicates). (D) Quantification of ‘free GFP’ protein level from (C) normalized to the Coomassie stained gel. Protein level in *PaSod3*^*H26L*^::*Gfp* cultures was set to 1. (E+F) Qualitative determination of superoxide anion and hydrogen peroxide release by histochemical NBT- and DAB-staining of *P. anserina* wild type and *ΔPaMic60, ΔPaMic19, ΔPaMic10* and *ΔPaMic26* strains. * Significant differences to the wild type \**p* < 0.05;\*\**p* < 0.01; \*\*\**p* < 0.001.

Similarly, in Mic10-subcomplex mutants we also observed a ‘free GFP’ increase. In *ΔPaMic10* as well as *ΔPaMic26* the amount of ‘free GFP’ significantly increases 1.5-fold compared to the wild type (Figure 5C,D), indicating an up-regulation of the mitophagy rate. In accordance with these data, in human cell lines and in *D. melanogaster* it was recently shown that mitophagy was induced after ablation of MIC60 (Guo et al., 2020; Wang et al., 2020). As expected, in the *PaMic12* deletion mutant no increase of ‘free GFP’ was detected (Figure S4). The rather slight increase of mitophagy in MICOS mutants is a marked difference from the strong increase of mitophagy (6-fold increase) in F_1_F_o_-ATP-synthase dimerization mutants (Warnsmann et al., 2021). The induction of mitophagy can have different causes. One of the most commonly described causes is the induction by oxidative stress due to increased levels of reactive oxygen species (ROS) (Graef & Nunnari, 2011; Lee et al., 2012). Accordingly, in HeLa cells missing MIC60 increased ROS production was found (John et al., 2005). To test whether or not ROS levels are altered in the *P. anserina* MICOS mutants, we used a histochemically method to visualize the extracellular amount of the two common ROS, superoxide anion and hydrogen peroxide (Munkres, 1990). This is an indirect measure of ROS levels in the different compartments of the cell. In contrast to the F_1_F_o_-ATP-synthase dimerization mutants (Warnsmann et al., 2021), we observed no changes in the hydrogen peroxide levels, but an increase in superoxide anion level in the MICOS mutants compared to the wild type (Figure 5E,F). The latter is indicated by the accumulation of a lilac precipitate specifically at the growth front of the mycelium. These data suggest that superoxide formation is the trigger for the mild mitophagy induction in MICOS mutants.

### 3.6 Lifespan extension of Mic10-subcomplex mutants is ROS-dependent

In biological systems, such as *Caenorhabditis elegans*, mild oxidative stress, due to elevated superoxide anion level, is known to cause longevity. This beneficial effect of mild oxidative stress is termed mitohormesis (Merry & Ristow, 2016; Ristow & Zarse, 2010). A similar lifespan extending adaptive response was described upon RNAi-mediated mitochondrial stress (Maglioni et al., 2014; Rea et al., 2007). In *P. anserina* it was also previously shown that superoxide anion-triggered mitophagy participates in mitohormesis (Knuppertz et al., 2017). Hence, we hypothesized that longevity of MICOS mutants is caused by mitohormesis. To experimentally validate this hypothesis, we analyzed the lifespan of the MICOS mutants and the wild type under oxidative stress induced by paraquat, a well-known herbicide which induces mitochondrial superoxide stress (Cocheme & Murphy, 2008). Normally, treatment with low paraquat concentrations (20-80 µM) results in a mitohormetic, pronounced lifespan extension, which is lost at higher paraquat concentrations (Knuppertz et al., 2017; Wiemer & Osiewacz, 2014). However, if the observed lifespan extension of the MICOS mutants is due to a ROS-dependent mitohormetic response, additional mild paraquat treatment should pass beneficial ROS levels and lead to a reduced lifespan. Consistent with previously published data (Knuppertz et al., 2017) we found that mild paraquat-induced oxidative stress has a lifespan-prolonging effect in the *P. anserina* wild type (Figure 6A,B; Table S6). The treatment with 80 µM paraquat prolongs the mean lifespan of the wild type by about 87% (43 d vs 23 d). Likewise, the lifespans of the *ΔPaMic60* and *ΔPaMic19* mutants are further increased by 80 µM paraquat compared to the untreated mutant (Figure 6A). The mean lifespans of the paraquat-treated *ΔPaMic60* and *ΔPaMic19* mutant are extended by at least 45% (68 d vs. 47 d) and 52% (76 d vs. 50 d), respectively (Figure 6B; Table S6). In contrast, the lifespans of the *ΔPaMic10* and *ΔPaMic26* mutants are decreased by treatment with 80 µM paraquat (Figure 6C,D; Table S7). The mean lifespans of the *ΔPaMic10* and *ΔPaMic26* mutants are reduced by at least 18% (32 d vs. 39 d) and 45% (29 d vs. 52 d), respectively (Figure 6D; Table S7). Thus, obviously 80 µM paraquat overwhelms the repair and recycling capacity of Mic10-subcomplex mutants, while Mic60-subcomplex mutants still have enough capacity. A mitohormetic response requires efficient transmission of the stress signal from the mitochondria to the cytosol and the nucleus. We speculate that this retrograde signaling is impaired upon ablation of Mic60-subcomplex due to its role in forming the contact between inner and outer mitochondrial membrane. Thus, although the appropriate signal (increased superoxide level) is present, the outcome is not identical in Mic10- and Mic60-subcomplex mutants. In Mic10-subcomplex mutants a strong mitohormetic response is induced in Mic10-subcomplex mutants, which is hardly effective in Mic60-subcomplex mutants. Additional ROS stress (80 µM paraquat) becomes detrimental in Mic10-subcomplex mutants leading to lifespan reduction. In contrast, in the Mic60-subcomplex mutants additional ROS stress (80 µM) is required to enhance the retrograde signal to stimulate a beneficial mitohormetic response. This conclusion suggests that in Mic60-subcomplex mutants a yet unknown pathway leads to longevity, while longevity in the Mic10-subcomplex mutants seems to be completely ROS-dependent.

**Figure 6:**
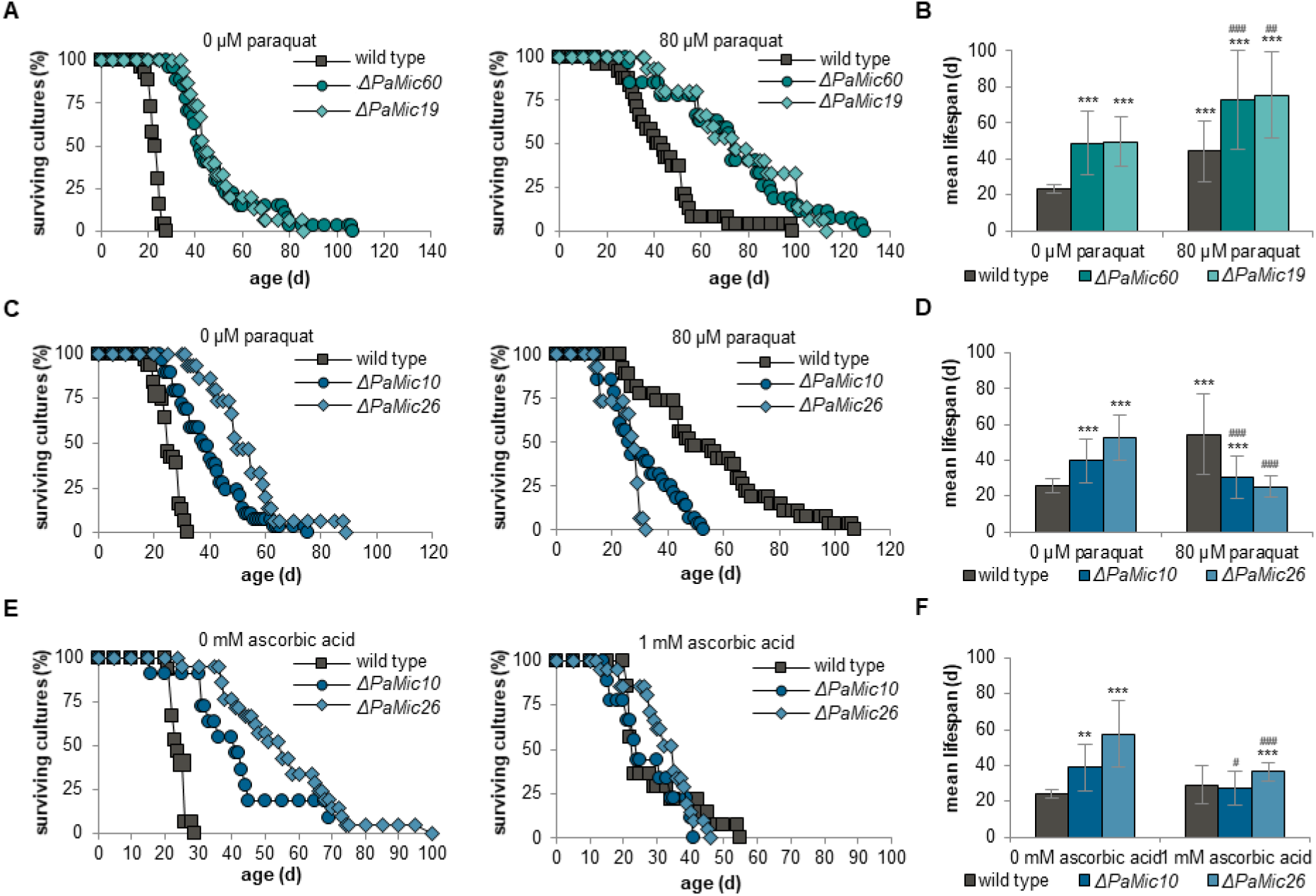
Lifespan extension of Mic10-subcomplex mutants is ROS-dependent. (A) Survival curves of *P. anserina* wild type (n = 26), *ΔPaMic60* (n = 27) and *ΔPaMic19* (n = 15) grown on standard M2 medium with or without 80 µM paraquat. (B) Mean lifespan of cultures from (A). (C) Survival curves of *P. anserina* wild type (n = 27), *ΔPaMic10* (n = 28) and *ΔPaMic26* (n = 27) grown on standard M2 medium with or without 80 µM paraquat. (D) Mean lifespan of cultures from (C). (E) Survival curves of *P. anserina* wild type (n = 15), *ΔPaMic10* (n = 11) and *ΔPaMic26* (n = 20) grown on standard M2 medium with or without 1 mM ascorbic acid. (F) Mean lifespan of cultures from (E). Data represent mean ± SD. * significant differences to the wild type without paraquat/ascorbic acid; ^#^ significant differences to the corresponding deletion mutant without paraquat/ascorbic acid. **^/##^*p* < 0.01; ***^/###^*p* < 0.001.

To verify this assumption, we analyzed the lifespan of *ΔPaMic10* and *ΔPaMic26* mutants under ROS scavenging conditions. Thereby the enhanced superoxide anions are scavenged by the antioxidant ascorbic acid. Under these conditions we obtained a reduction of the *ΔPaMic10* and *ΔPaMic26* lifespan (Figure 6E,F; Table 8). The mean lifespan of the *ΔPaMic26* mutant is reduced by more than 36% (37 d vs. 58 d) compared to the untreated mutant (Figure 6F; Table 8). Comparably, the mean lifespan of the *ΔPaMic10* mutant is reduced by more than 35% (27 d vs. 36 d). These data verify that longevity of Mic10-subcomplex mutants depends on ROS-induced mitohormesis.

Retrograde signals from mitochondria upon oxidative stress and damage can be very diverse. For instance, peptides from the proteolytic degradation of damaged or misfolded proteins, a reduced mitochondrial membrane potential or lipid peroxidation and a consequently altered lipid composition of the outer mitochondrial membrane can activate signal cascades. Moreover, a congestion of protein transport into mitochondria can act as a stress signal (Callegari & Dennerlein, 2018; Hill & Van Remmen, 2014; Kim et al., 2016; Tan & Finkel, 2020). As mentioned above, we suggest that retrograde signaling is impaired in Mic60-subcomplex mutants because of the compromised contact between inner and outer mitochondrial membrane. This hypothesis is confirmed by the observation that mitochondrial import defects are consequences of MICOS disruption in yeast (Khosravi & Harner, 2020; Wiedemann & Pfanner, 2017). For an efficient protein import, an interaction between the Mic60-subcomplex and the protein import machinery is necessary (Harner et al., 2011; Körner et al., 2012; von der Malsburg et al., 2011).

### 3.7 Both MICOS subcomplexes act in parallel pathways to regulate lifespan

If the mitohormetic response is attenuated in the Mic60-subcomplex mutants and in Mic10-subcomplexes this response is active leading to lifespan extension, a double mutant lacking both subcomplexes should show an additive effect and further lifespan extension. To test this possibility, we generated double mutants in which one subunit of each subcomplex is missing (Figure S5). And indeed, a double mutant missing both core subunits shows a spectacular lifespan extension compared to the single mutants (Figure 7A,B). The mean lifespan of the *ΔPaMic60/ΔPaMic10* mutant is increased by 100% and 159%, respectively, compared to the single *ΔPaMic60* and *ΔPaMic10* mutants (Figure 7B; Table S8). Likewise, the simultaneous deletion of *PaMic60* and *PaMic26* was also found to be synergistic leading to mean lifespan extension to more than 121 days (Figure 7C,D; Table S8). This is an increase of 142% and 133%, respectively, compared to the *ΔPaMic60* and *ΔPaMic26* single mutants. The lifespan analyses of the double mutants revealed that the longevity of Mic60-subcomplex mutants is based on a pathway that is distinct from that of the Mic10-subcomplex mutants.

**Figure 7:**
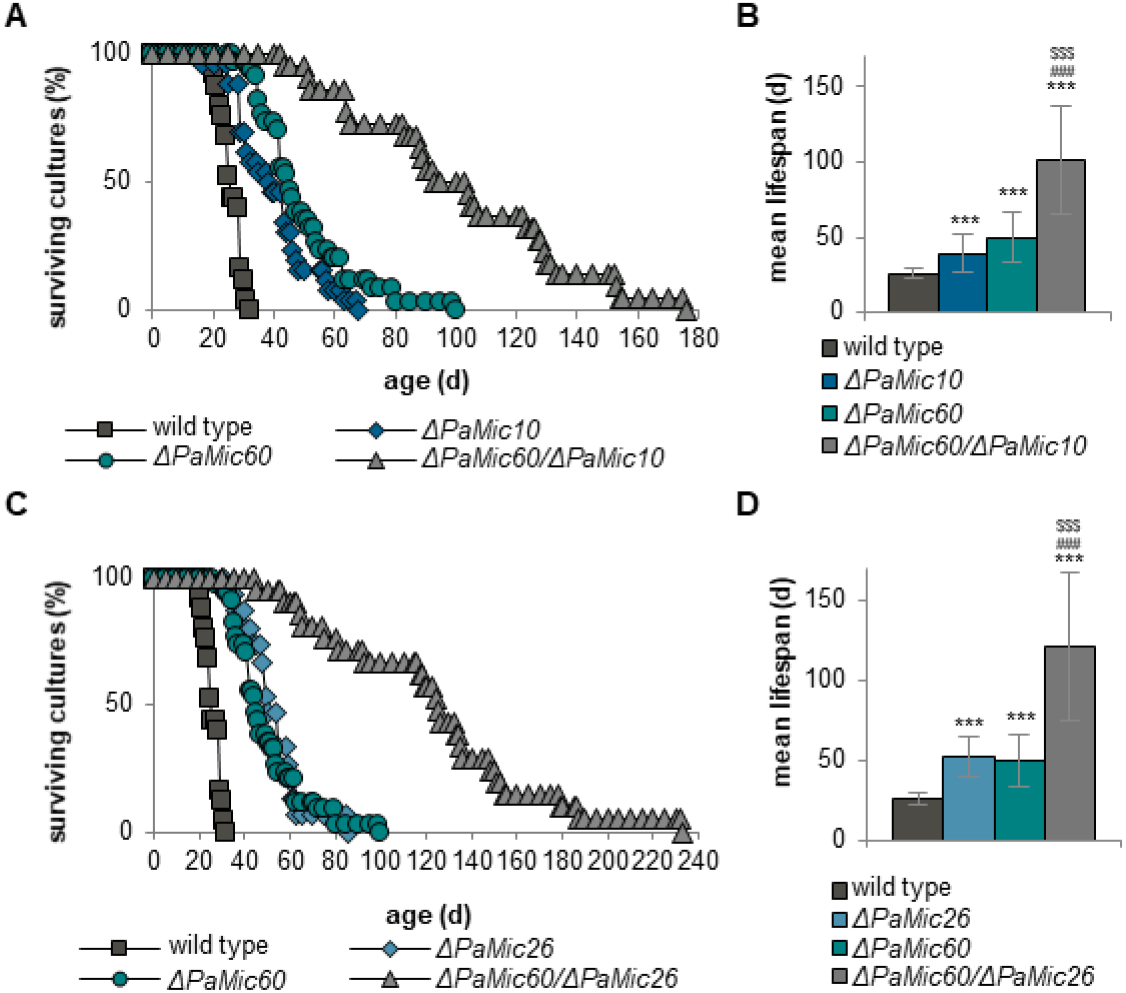
Simultaneous ablation of subunits from both MICOS subcomplex synergistically increases lifespan. (A) Survival curves of *P. anserina* wild type (n = 25), *ΔPaMic60* (n = 34), *ΔPaMic10* (n = 27) and *ΔPaMic60/ΔPaMic10* (n = 22) grown on standard M2 medium. (B) Mean lifespan of cultures from (A). (C) Survival curves of *P. anserina* wild type (n = 25), *ΔPaMic60* (n = 34), *ΔPaMic26* (n = 15) and *ΔPaMic60/ΔPaMic26* (n = 23) grown on standard M2 medium. (D) Mean lifespan of cultures from (C). Data represent mean ± SD. *** significant differences to wild type; ^###^ significant differences to *ΔPaMic26* or *ΔPaMic10;* ^$$$^ significant differences to *ΔPaMic60. p* < 0.001.

Previous studies point out a connection between the MICOS complex and mitochondrial phospholipids. In yeast, the MICOS complex is necessary for phospholipid transfer between inner and outer mitochondrial membrane (Aaltonen et al., 2016). Moreover, in human cells proteins of the phospholipid metabolism, e.g. the cardiolipin synthase, interact in a large protein complex with mitochondrial membrane proteins like MIC60 and MIC19 (Serricchio et al., 2018). It is thus well possible that MICOS alteration, specifically Mic60-subcomplex ablation affects phospholipid homeostasis. Since it was recently shown that in *P. anserina* mitochondrial phospholipid homeostasis affects the aging process (Löser et al., 2021), longevity of Mic60-subcomplex mutants may be related to the mitochondrial phospholipid homeostasis. In fact, preliminary studies show that *PaMic60* deletion leads to an enhanced abundance of the cardiolipin synthase PaCRD1 (data not shown). However, further studies are required to clarify the relationship between phospholipid metabolism and lifespan of Mic60-subcomplex mutants.

## 4 CONCLUSIONS

In the current study we uncovered a counterintuitive and unexpected biological role of MICOS. Ablation of MICOS components in *P. anserina* leads to changes in mitochondrial cristae ultrastructure and morphology and to a pronounced lifespan extension. By the use of double mutants lacking both subcomplexes, we provide first genetic evidence that the two MICOS subcomplexes impact organismic aging via different pathways. While longevity of Mic10-subcomplex mutants is triggered by ROS-induced mitohormesis, the pathway leading to longevity of the Mic60-subcomplex mutants is yet unclear but may be linked to phospholipid homeostasis. This distinction may be due to the role of MIC60 in forming the contact between outer and inner mitochondrial membrane as well as in mitochondrial protein import. Future studies need to address the question whether this longevity pathway in MICOS mutants is evolutionary conserved from simple organisms like *P. anserina* up to humans.

## ACKNOWLEDGEMENTS

We are grateful to Dr. Andrea Hamann for data discussion.

This work was funded by the Deutsche Forschungsgemeinschaft (DFG, German Research Foundation) –Project-ID 25913077–SFB1177 to HDO and SE and Os75/17-2 to HDO.

## CONFLICT OF INTEREST

None declared.

## AUTHOR CONTRIBUTIONS

VW and HDO designed this study and wrote the manuscript. VW, LMM, ACM, MW, LS, MB and SE performed experiments. VW, LMM, SE and HDO analyzed data. VW visualized data. HDO, SE and VW reviewed and edited the manuscript. HDO supervised the study and acquired funding. All authors have read the final version of the manuscript.

## DATA AVAILABILITY STATEMENT

The data that support the findings of this study are available on request from the corresponding author.

## Notes

### Competing Interest Statement

The authors have declared no competing interest.

