## Supplements for "Disruption of the MICOS complex leads to an aberrant cristae structure and an unexpected, pronounced lifespan extension in *Podospora anserina*"

#### Figure S1

**A**

```
PaMIC60       MLRTSIRSVRALGSRPAAAVAGRQWQASAARRAVVSGQRFYADDKKPVVDEAKLPAAETLTAPTTPPPPPPQTTPAATITPEQAPLTPPAPGP
ScMIC60       ---------------------------------------------------------------------------------------------

AVVPPPPPPVAPIVPKKKGFFRR-------------------LRNFLLKLVLLGVLGFGGGVWYSRINDNFHDFFTQYVPYGEEAV-----------LYLEELDFRK
--------MMLRTTASRKIVLRRGLASINTGTTVASKKASHKFRNTLWTIALSATAFYAGGIIYSQKNDKFGDFFSNNVPFAEDLLETYEHYHDRPTLFLED-----

RFPNVANRVTGRRADTASEQVSIPAQSGASWRVADGDHAGRQSSGVAQKVVAAKDAVKDTVKDIVKPEPAAVTKAKKETAALPKVEAKDAALEKKVKGVEEKKEVAV
-----------------------------SW-----------------------DGLKAKSNDL-----------------LSGLTGSSQTRRSNRENIEVKKILS-

IPAAATSSKPKEFRAPDVNEPSVWPPASPIDPLSV-PHADDAIVQQLVHMLNDIITVINYDGAADKYSQTIWKAKDEVSKVGERILSIKSAAEEEAAKQVKARIDSF
-----------------------------LEPLNIETENSDPQLKEIIGSLNDLINSLNDSNLSIPESEF-----NSIKKSNQNMLTNLSQLNETLKEAL-------

DKHANDLVSRLEAIMLAQEQQW---RQEFEAEVER-LKHNYDD-KIKLIQEREQKLSEQKLQNKLLEQA----------VELQRQFSRDIKKHVEEEREGRLGRIES
---SNYMIQRTSEVITELNTQYENSKREFEKNLQKNLLQEVDEFKENLTKQKDKELEEKLKANEELLQAKHANEVGLLSITQVKEFNKIIKDKIEKERNGRLAHLEE

LSRAVSELEKLTTGLNEVVDTNLRTQQLHVAVEAVRASLEDAHHPRPFI-KELVALKEIA-----------ADD------------------------------PVV
INSEVNDLSKSIDRSSKILSKNEALVQLTFQVDEIKSRINNNNLPDVNIDKELSRLKLLSNLLSTFNKKSCCDDGDCCSCKKGNKNEGKEGKISCKCKPKTNPPSLL

DAAIASIHPSAYQRGISTSAELIDRFRRVAAEVRKASLLPEDAGVASHASSYLLSKVMFKKQGLAA-GDDVESILTRTQTFLEEGDLDNAAREMNTLGGWAKTLSRD
SVALDELESTCSGKKILSNEQIYNRWNLLADDFKTASLLPPNSGILGQLTAKVFSLFLFTKTGNPSNATDFDSVYARVGDNLRVSNLNDAVEEVVSLKGWPHKVCES

WLSEVRKVLE VQQALDVITTEARLQSLKVE    676 aa
WIEDARRKLE VQRLVEILDCEIRTL-----    540 aa
```

**B**

```
PaMIC19       MGSSSSKPAATA--PQTWKAPGSAGLSTELVQHLHSSPETDASRLQAIELEIQARVAAELKRLRDQESEALRAAQAKLAE---STDDKTADDN
ScMIC19       MGSNTSKVGAGAEKQQVYTPLTQIDFSQSLVSQLDSSKESDYVTKQNAEKFIEKKVS---QRLSNLEVETLKKFEDTLNNSLLSDDDKDAVDG

SKTSYTVSKEVQALQKKLEERKRIRSQLPESLENA---------------RSGVVRCLRENDRRPLDCWREVEAFKEEVRRVEKGWVEKVVS  165 aa
ISSS-SLNNQIESLNKKLTLFDQLELQKLEKYGGAKGKSDKKTDNGSISIKAKLTECLLANKGKPLNCYEEMEEFKKLVMG-----------  170 aa
```

**C**

```
PaMIC10       ---------MADTAPATTTSPTSRAVSRPVSEALLNEKWDRCFSNLIIKSSLGLGFGVVFSVLLFKRRAWPAFVGVGFGAGRAYEECNSSLKQ
ScMIC10       MSEQAQTQQPAKSTPSKDSNKNGSSVS-----TILDTKWDIVLSNMLVKTAMGFGVGVFTSVLFFKRRAFPVWLGIGFGVGRGYAEGDAIFRS

AAKEIRKQA-     93 aa
SA-GLRSSKV     97 aa
```

**D**

```
PaMIC26       MAARLLFQRRAAPLTAAVLVGSIAFSPSLAHAEAPNGRQFPRKPIYDDDLDDPLPTSKTLPAPPKPAAAEPTTPSTSLITLPTTSEPAVIKKE
ScMIC26       -------------------------------------------------------MTKDFYRQLDPVEEKIVPPENAIVISSEAKEATVNEKE

TPTDLLAQRIRSARLYLYHHSCI----------------------------LEDAVNSSLSRAFDLEQSFTSTISGLAPDRASGEKLMPGAIYVLVAGMAGSIVARN
AKQGVLSQRVMK---YIGENELVDGISVRDPDYLKRFFNERRKQFSAKWDKVTNKIDDIAGRYYAREESFTSTIASLHTD--PNERLIPGLLSILVASMTGSVLARR

RNVLLRTATPLALGITAAWTVLPVTMTNVSELLWYYEKKV-PAVAEAHLQVREGVEKGVYMAKVHAELAQRKIEEGVREVREGLEGW----VRKGK*    264 aa
RTWLLRATMPIILGSCCFAYAMPTTFRNTMGLIHNLEMNTFPHFTERQDRVWKETKRLSTASVQYYYDAKKWLNKDVEKTGNAIKNWTGVNVK----    233 aa
```

**E**

```
PaMIC12       --------------------------MRFSWGRTGGLTLTLSLTYLALHTHRRNREVQSTLLRSQSHTLDTLNPLSSSTLPPRKSSYITTDGT
ScMIC12       MSKLGPLARSVKWTLSVGVIGSVFYLYRYS---NNGYFYDHDATWL-----KQDHQVQDLVDRKEVVPGETRN----------RKLVVTDDGT

HTYIPRQTLASTTTSFLEEAKSRWNSEVIAAVNWAQSHSPAIESTASSAVVDLAHNAPAIKERVADASREGWQTAEKGLFRAEKKIGEVVERTVEKGKEVYGKAKAK
----------AWSRTMGESIKDIWNEQIRNSVDWIYSWGKN------------------------------------------------------------------

VYLAEEKLEGKLEAKLEGVSEIERALAERFDQAKRERRLERSVEEVLAERYKPIDERDFGGLALI     239 aa
-----------------------------------------------------------------     106 aa
```

**Figure S1**: Amino acid sequence alignments of *P. anserina* MICOS proteins with their yeast homologs. Amino acid sequence alignment of (A) *P. anserina* MIC60 (PaMIC60; UniProt: B2A9R4) and *S. cerevisiae* MIC60 (ScMIC60; UniProt: P36112) (B) *P. anserina* MIC19 (PaMIC19; UniProt: B2AUM5) and *S. cerevisiae* MIC19 (ScMIC19; UniProt: P43594), (C) of *P. anserina* MIC10 (PaMIC10; UniProt: B2AWQ3) and *S. cerevisiae* MIC10 (ScMIC10; UniProt: Q96VH5), (D) of *P. anserina* MIC26 (PaMIC26; UniProt: B2AYB9) and *S. cerevisiae* MIC26 (ScMIC26; UniProt: P50087) and (E) of *P. anserina* MIC12 (PaMIC12; UniProt: B2B4I6) and *S. cerevisiae* MIC12 (ScMIC12; UniProt: P38341) using EMBOSS Needle (http://www.ebi.ac.uk/Tools/psa/emboss_needle/). Identical amino acids are marked by grey boxes. Conserved domains are marked by blue lines in the *S. cerevisiae* sequences.

### Figure S2

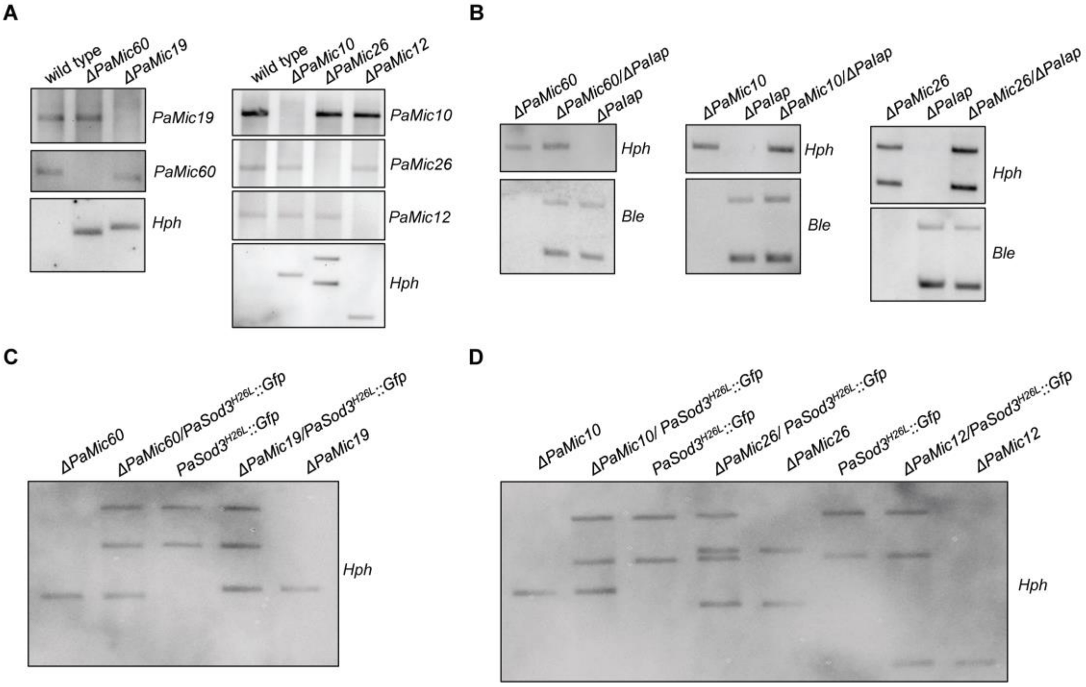


**Figure S2:** Southern blot verification of MICOS mutants. (A) Southern blot analyses verifying the replacement of *PaMic60*, *PaMic19*, *PaMic10*, *PaMic26* and *PaMic12* by a hygromycin resistance gene (*Hph*). *PaMic60*, *PaMic19*, *PaMic10*, *PaMic26*, *PaMic12* and *Hph* probes were used as indicated. (B) Southern blot analyses verifying the newly generated double mutants *ΔPaMic60/ΔPaIap*, *ΔPaMic10/ΔPaIap* and *ΔPaMic26/ΔPaIap*. *Ble* (phleomycin resistance), *Hph* (hygromycin resistance) probes were used as indicated. (C+D) Southern blot analysis validating the genetic constitution of different *PaSod3*$^{H26L}$*::Gfp* mutants and their respective control strains. *Hph* (hygromycin resistance) probe was used.

### Figure S3

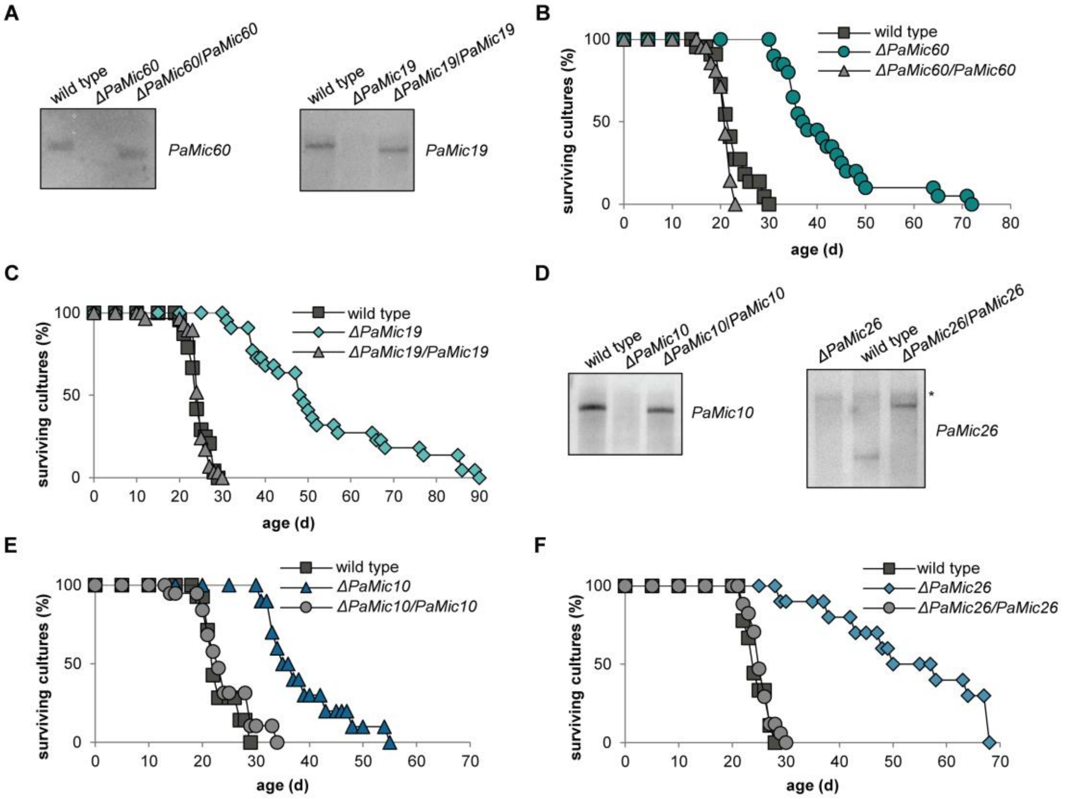


**Figure S3:** Southern blot verification and lifespan analysis of MICOS complementation strains. (A) Southern blot analysis validates the genotype of different complementation strains and their respective control strains. *PaMic60* and *PaMic19* specific probes were used to detect the corresponding genes. (B) Survival curves of *P. anserina* wild type (n = 22); *ΔPaMic60* (n = 20) and *ΔPaMic60/PaMic60* (n = 21) grown on standard M2 medium. (C) Survival curves of *P. anserina* wild type (n = 24); *ΔPaMic19* (n = 22) and *ΔPaMic19/PaMic19* (n = 29) grown on standard M2 medium. (D) Southern blot analysis validates the genotype of different complementation strains and their respective control strains. *PaMic10* and *PaMic26* specific probes were used to detect the corresponding genes. An asterisk (*) marks an unspecific signal. (E) Survival curves of *P. anserina* wild type (n = 14); *ΔPaMic10* (n = 10) and *ΔPaMic10/PaMic10* (n = 19) grown on standard M2 medium. (F) Survival curves of *P. anserina* wild type (n = 9); *ΔPaMic26* (n = 10) and *ΔPaMic26/PaMic26* (n = 17) grown on standard M2 medium.

### Figure S4

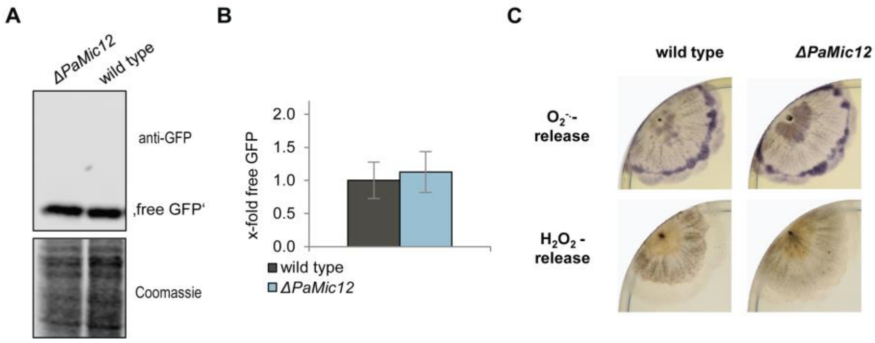


**Figure S4:** Mitophagy and ROS release in the *PaMic12* deletion mutant. (A) Monitoring mitophagy by western blot analysis of total protein extracts from *PaSod3*$^{H26L}$*::Gfp* (here wild type) and *ΔPaMic12/PaSod3*$^{H26L}$*::Gfp* (*ΔPaMic12*) cultures with a GFP antibody (each 4 biological replicates). (B) Quantification of 'free GFP' protein level normalized to the Coomassie stained gel. Protein level in *PaSod3*$^{H26L}$*::Gfp* cultures was set to 1. Data represent mean ± SD. (C) Qualitative determination of superoxide anion and hydrogen peroxide release, respectively, by histochemical NBT- and DAB-staining of *P. anserina* wild-type and *ΔPaMic12* strains.

#### Figure S5

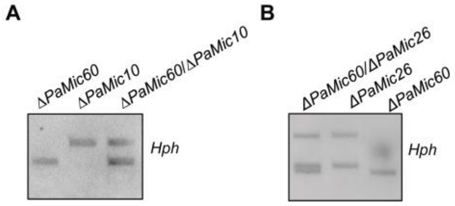


**Figure S5:** Southern blot analysis of MICOS double deletion mutants. Southern blot analysis verifies the genotype of (A) *ΔPaMic60/ΔPaMic10* and of (B) *ΔPaMic60/ΔPaMic26* and their respective control strains using an *Hph*-specific gene probe.

#### Table S1

**Table S1:** Oligonucleotides used for amplification of flanking regions for deletion of specific genes. The restriction sites are underlined in the sequences. All oligonucleotides were purchased from Biomers, Ulm, Germany.

| **Gene** | **oligonucleotide** | | **restriction site** |
|---|---|---|---|
| | **name** | **sequence (5'-3')** | |
| *PaMic10 (Pa_7_7950)* | PaMic10_KO5 | GCTCTGCAGATCACCAAATCAAATTCACGC | PstI |
| | PaMic10_KO6 | GAGGATCCTTGCTGTCTTCTTCTACC | BamHI |
| | Mic10OEx1 | CAGGTACCGCTAGGGAGAAGAGAGG | KpnI |
| | Mic10OEx2 | GCGGAAGCTTGTGATTGTGTGGGAGTG | HindIII |
| *PaMic12 (Pa_2_1415)* | PaMic12-KO1 | GGATCGATGACCGCCTTGTCCTTGAG | ClaI |
| | PaMic12-KO2 | GCCGAAGCTTGTTGTTCAGATAACCGTTG | HindIII |
| | PaMIc12-KO3 | GCACTGCAGGGGAGGTTGATAAGGGGAG | PstI |
| | PaMic12-KO4 | GAGGATCCGATTGGGCCGTTGATTGCG | BamHI |
| *PaMic26 (Pa_1_10590)* | PaMic26-KO1 | GCATCGATACCGCTGGGCTACCTAGAGC | ClaI |
| | PaMic26-KO2 | GCCGAAGCTTGGTCTCGACCCAATTGG | HindIII |
| | PaMic26-KO3 | GCTCTGCAGGGGGAATGGTTGTGTG | PstI |
| | PaMic26-KO4 | GAGGATCCCTACCGCCAAACCACTCTC | BamHI |
| *PaMic60 (Pa_1_1530)* | PaMic60-KO1 | GAGGATCCAGGACTCAGGCGTATGAC | BamHI |
| | PaMic60-KO2 | GACTGCAGGAGGGAGATGGAGTGGCC | PstI |
| | PaMic60-KO3 | GCGATATCAAGGGACCAACGGGACGTC | EcoRV |
| | PaMic60-KO4 | GCAAGCTTTGCCCTTGCCTTTTGTCC | HindIII |
| *PaMic19 (Pa_1_19620)* | PaMic19-KO1 | GCGATCGATGTTGTACGAGCATCAAG | ClaI |
| | PaMic19-KO2 | GGAAGCTTGTGAGGGATTGTCTGGTTG | HindIII |
| | PaMic19-KO3 | GGCTGCAGGGTGGTAACGAAGTTGG | PstI |
| | PaMic19-KO4 | GCGGATCCATGATCCCATCTCTTC | BamHI |

#### Table S2

**Table S2:** Oligonucleotides used for amplification of genes for complementation. The restriction sites are underlined in the sequences. All oligonucleotides were purchased from Biomers, Ulm, Germany.

| **Gene** | **oligonucleotide** | | **restriction site** |
|---|---|---|---|
| | name | sequence (5'-3') | |
| *PaMic10 (Pa_7_7950)* | Mic10OEx1 | CAGGTACCGCTAGGGAGAAGAGAGG | KpnI |
| | Mic10-Kom1 | GTGTCGACGTGGATTTAGGCCGAGGATG | SalI |
| *PaMic26 (Pa_1_10590)* | Mic26-3 | GAGTCGACGCCACCTATGTCGATAAC | SalI |
| | Mic26-4 | CGATCGATGAGGGAGATTGTAGCG | ClaI |
| *PaMic60 (Pa_1_1530)* | PaMic60_1 | GTCTCGAGTATGACCGGCAGTGGATG | XhoI |
| | PaMic60_2 | GAAGAGCCGCAGTCTCCTTC | 5'Phosphat |
| | PaMic60_3 | CCAAGGTCGAAGCCAAGGATG | 5'Phosphat |
| | PaMic60_4 | CAGGTACCGGACCGGAGTAAGACTG | KpnI |
| *PaMic19 (Pa_1_19620)* | PaMic19_KO1 | GCGATCGATGTTGTACGAGCATCAAG | ClaI |
| | PaMic19-5 | GAGGTACCGGAGGTTTAGGGAGGCACTG | KpnI |

#### Table S3

**Table S3:** Oligonucleotides used for the construction of specific gene probes. All oligonucleotides were purchased from Biomers, Ulm, Germany.

| Gene | oligonucleotide | | base pairs (bp) |
|---|---|---|---|
| | **name** | **sequence (5'-3')** | |
| *PaMic10 (Pa_7_7950)* | Pa_7_7950-1 | TCAGCGAAGCCCTCCTCAAC | 311 |
| | Pa_7_7950-2 | TAGGCTTGCTTCCTGATCTC | |
| *PaMic12 (Pa_2_1415)* | PaMic12-1 | CTAACCTACCTAGCCCTCC | 289 |
| | PaMic12-2 | GCGGTTGATTCGATAGCG | |
| *PaMic26 (Pa_1_10590)* | PaMic26-1 | GGTCGTCAGTTTGTACGTC | 297 |
| | PaMIc26-2 | GATGCAGGAGTGGTGGTAG | |
| *PaMic60 (Pa_1_1530)* | Pa_1_1530-1 | GCGTACATCGATACGATCAG | 319 |
| | Pa_1_1530-2 | GTGCTTGCTCGGGAGTAATC | |
| *PaMic19 (Pa_1_19620)* | PaMic19-1 | CCAGCACCTCCATTCCTC | 306 |
| | PaMic19-2 | CTTCTTCTGGAGAGCTTGC | |

#### Table S4

**Table S4:** Oligonucleotides used for the quantitative real-time PCR (qPCR). All oligonucleotides were purchased from Biomers, Ulm, Germany.

| Gene | oligonucleotide | |
|---|---|---|
| | **name** | **sequence (5'-3')** |
| *PaMic10 (Pa_7_7950)* | Pa_7_7950-1 | TCAGCGAAGCCCTCCTCAAC |
| | Pa_7_7950-2 | TAGGCTTGCTTCCTGATCTC |
| *PaMic60 (Pa_1_1530)* | Pa_1_1530-3 | GCCTCTAGCTACCTCTTG |
| | Pa_1_1530-4 | TTGCAGTCTGGCCTCGGTTG |
| *PaMic19 (Pa_1_19620)* | PaMic19-1 | CCAGCACCTCCATTCCTC |
| | PaMic19-2 | CTTCTTCTGGAGAGCTTGC |
| *PaPorin (Pa_2_9780)* | Porin-RT-for | TCTCCTCCGGCAGCCTTG |
| | Porin-RT-rev | CGGAGGCGGACTTGTGAC |

#### Table S5

**Table S5:** Overview of lifespan and growth rate of *P. anserina* wild type and MICOS deletion mutants. The *p*-values were determined with SPSS (IBM; SPSS Statistics, New York, USA) with three different tests. Indicated are the *p*-values of the lifespan curves in comparison to the wild type. "*p*-value 1" = Log Rank (Mantel-Cox); "*p*-value 2" = Breslow (Generalized Wilcoxon); "*p*-value 3" = Tarone-Ware.

| | wild type | *ΔPaMic60* | *ΔPaMic19* | *ΔPaMic10* | *ΔPaMic12* | *ΔPaMic26* |
|---|---|---|---|---|---|---|
| mean lifespan (d) ± SD | 25 ±3.97 | 47 ±16.37 | 50 ±16.05 | | 23 ±3.75 | 52 ±14.96 |
| Maximum lifespan (d) | 32 | 105 | 90 | | 33 | 100 |
| *p*-value 1 | / | 3.29E-37 | 1.84E-25 | 2.21E-18 | 0.13 | 7.87E-33 |
| *p*-value 2 | / | 9.39E-32 | 1.93E-20 | 3.97E-15 | 0.06 | 1.28E-27 |
| *p*-value 3 | / | 1.76E-34 | 6.28E-23 | 9.27E-17 | 0.07 | 3.33E-30 |
| growth rate (cm/d) ± SD | 0.67 ±0.05 | 0.62 ±0.04 | 0.62 ±0.03 | 0.62 ±0.04 | 0.65 ±0.09 | 0.65 ±0.03 |
| growth distance (cm)± SD | 14.2 ±2.6 | 27.5 ±9.7 | 30.3 ±9.8 | | 13.9 ±2.1 | 30.7 ±9.5 |
| biological replicates | 66 | 74 | 45 | 48 | 54 | 66 |

### Table S6

**Table S6:** Overview of lifespan and growth rate of *P. anserina* wild type and Mic60-subcomplex deletion mutants with and without paraquat. The *p*-values were determined with SPSS (IBM; SPSS Statistics, New York, USA) with three different tests. Indicated are the *p*-values of the lifespan curves in comparison to the wild type without paraquat. "*p*-value 1" = Log Rank (Mantel-Cox); "*p*-value 2" = Breslow (Generalized Wilcoxon); "*p*-value 3" = Tarone-Ware. Additionally, *p*-values of the lifespan curves compared to the corresponding deletion mutant without paraquat are given. "*p*-value 4" = Log Rank (Mantel-Cox); "*p*-value 5" = Breslow (Generalized Wilcoxon); "*p*-value 6" = Tarone-Ware.

| | wild type | *ΔPaMic60* | *ΔPaMic19* | wild type | *ΔPaMic60* | *ΔPaMic19* |
|---|---|---|---|---|---|---|
| | 0 µM paraquat | | | 80 µM paraquat | | |
| mean lifespan (d) ± SD | 23 ±2.4 | 48 ±17.8 | 50 ±13.8 | 44 ±16.8 | 72 ±27.5 | 76 ±24.3 |
| Maximum lifespan (d) | 28 | 107 | 86 | 99 | 129 | 113 |
| *p*-value 1 | / | 7.96E-15 | 1.03E-09 | 3.67E-11 | 7.96E-15 | 1.03E-09 |
| *p*-value 2 | / | 1.18E-12 | 8.09E-08 | 4.24E-09 | 1.18E-12 | 8.09E-08 |
| *p*-value 3 | / | 1.00E-13 | 9.56E-09 | 4.13E-10 | 1.00E-13 | 9.56E-09 |
| *p*-value 4 | / | / | / | / | 0.0003 | 0.001 |
| *p*-value 5 | / | / | / | / | 0.001 | 0.003 |
| *p*-value 6 | / | / | / | / | 0.0004 | 0.002 |
| growth rate (cm/d) ± SD | 0.62 ±0.04 | 0.60 ±0.04 | 0.57 ±0.02 | 0.59 ±0.04 | 0.55 ±0.03 | 0.57 ±0.02 |
| growth distance (cm) ± SD | 13.2 ±1.8 | 27.4 ±10.9 | 29.7 ±7.5 | 21.4 ±8 | 32.5 ±11.1 | 36.1 ±12.2 |
| biological replicates | 26 | 27 | 15 | 26 | 27 | 15 |

#### Table S7

**Table S7:** Overview of lifespan and growth rate of *P. anserina* wild type and Mic10-subcomplex deletion mutants with and without paraquat or ascorbic acid. The *p*-values were determined with SPSS (IBM; SPSS Statistics, New York, USA) with three different tests. Indicated are the *p*-values of the lifespan curves in comparison to the wild type without paraquat or ascorbic acid. "*p*-value 1" = Log Rank (Mantel-Cox); "*p*-value 2" = Breslow (Generalized Wilcoxon); "*p*-value 3" = Tarone-Ware. Additionally, *p*-values of the lifespan curves compared to the corresponding deletion mutant without paraquat or ascorbic acid are given. "*p*-value 4" = Log Rank (Mantel-Cox); "*p*-value 5" = Breslow (Generalized Wilcoxon); "*p*-value 6" = Tarone-Ware.

| | wild type | *ΔPaMic10* | *ΔPaMic26* | wild type | *ΔPaMic10* | *ΔPaMic26* |
|---|---|---|---|---|---|---|
| | 0 µM Paraquat | | | 80 µM Paraquat | | |
| mean lifespan (d) ± SD | 26 ±4.1 | 40 ±12.4 | 53 ±12.6 | 55 ±22.6 | 30 ±11.6 | 25 ±6.1 |
| Maximum lifespan (d) | 32 | 75 | 89 | 107 | 53 | 32 |
| *p*-value 1 | / | 5.65E-09 | 7.24E-10 | 9.91E-10 | 0.02 | 0.75 |
| *p*-value 2 | / | 2.61E-07 | 7.90E-08 | 8.53E-08 | 0.31 | 0.76 |
| *p*-value 3 | / | 4.11E-08 | 7.86E-09 | 1.02E-08 | 0.10 | 0.72 |
| *p*-value 4 | / | / | / | / | 0.01 | 1.87E-08 |
| *p*-value 5 | / | / | / | / | 0.004 | 1.99E-07 |
| *p*-value 6 | / | / | / | / | 0.007 | 5.96E-08 |
| growth rate (cm/d) ± SD | 0.68 ±0.03 | 0.61 ±0.04 | 0.66 ±0.01 | 0.59 ±0.1 | 0.54 ±0.03 | 0.51 ±0.02 |
| growth distance (cm) ± SD | 14.8 ±2.6 | 20.4 ±5.8 | 30.8 ±7.7 | 25.6 ±11.2 | 14.2 ±7.3 | 10.6 ±1.8 |
| biological replicates | 27 | 28 | 15 | 27 | 28 | 15 |
| | 0 mM ascorbic acid | | | 1 mM ascorbic acid | | |
| mean lifespan (d) ± SD | 24 ±2.2 | 39 ±12.9 | 58 ±18.5 | 29 ±10.8 | 27 ±9.1 | 37 ±5.3 |
| Maximum lifespan (d) | 29 | | 100 | 55 | 41 | 46 |
| *p*-value 1 | / | 2.01E-05 | 1.12E-09 | 0.322 | 0.173 | 7.26E-06 |
| *p*-value 2 | / | 2.67E-04 | 9.48E-09 | 0.982 | 0.691 | 1.22E-04 |
| *p*-value 3 | / | 7.55E-05 | 3.22E-09 | 0.661 | 0.402 | 2.98E-05 |
| *p*-value 4 | / | / | / | / | 0.01 | 3.39E-07 |
| *p*-value 5 | / | / | / | / | 0.02 | 4.37E-06 |
| *p*-value 6 | / | / | / | / | 0.01 | 1.23E-06 |
| growth rate (cm/d) ± SD | 0.65 ±0.03 | 0.67 ±0.02 | 0.64 ±0.01 | 0.61 ±0.01 | 0.60 ±0.04 | 0.60 ±0.01 |
| growth distance (cm) ± SD | 14.1 ±1.4 | | 35.3 ±12.4 | 16.1 ±6.8 | 15.7 ±5.4 | 20.1 ±2.9 |
| biological replicates | 15 | 11 | 21 | 15 | 11 | 21 |

#### Table S8

**Table S8:** Overview of lifespan and growth rate of *P. anserina* wild type and MICOS deletion mutants and double deletion mutants**.** The *p*-values were determined with SPSS (IBM; SPSS Statistics, New York, USA) with three different tests. Indicated are the *p*-values of the lifespan curves in comparison to the wild type. “*p*-value 1” = Log Rank (Mantel-Cox); “*p*-value 2” = Breslow (Generalized Wilcoxon); “*p*-value 3” = Tarone-Ware. Moreover, *p*-values of the lifespan curves compared to *ΔPaMic10* or *PaMic26*. “*p*-value 4” = Log Rank (Mantel-Cox); “*p*-value 5” = Breslow (Generalized Wilcoxon); “*p*-value 6” = Tarone-Ware. And, *p*-values of the lifespan curves compared to *ΔPaMic60*. “*p*-value 7” = Log Rank (Mantel-Cox); “*p*-value 8” = Breslow (Generalized Wilcoxon); “*p*-value 9” = Tarone-Ware.

| | wild type | *ΔPaMic60* | *ΔPaMic10* | *ΔPaMic26* | *ΔPaMic60/ ΔPaMic10* | *ΔPaMic60/ ΔPaMic26* |
|---|---|---|---|---|---|---|
| mean lifespan (d) ± SD | 26 ±3.5 | 50 ±16.1 | 39 ±12.5 | 52 ±12.6 | 101 ±35.8 | 121 ±45.9 |
| Maximum lifespan (d) | 32 | 100 | 63 | 86 | 176 | 233 |
| *p*-value 1 | / | 2.82E-16 | 2.23E-07 | 1.70E-09 | 1.17E-12 | 3.09E-12 |
| *p*-value 2 | / | 2.52E-14 | 4.26E-06 | 9.63E-08 | 1.22E-10 | 3.09E-10 |
| *p*-value 3 | / | 2.72E-15 | 1.04E-06 | 1.33E-08 | 1.26E-11 | 3.27E-11 |
| *p*-value 4 | / | 0.02 / 0.59 | / | / | 8.88E-11 | 2.64E-08 |
| *p*-value 5 | / | 0.01 / 0.24 | / | / | 4.19E-09 | 2.61E-07 |
| *p*-value 6 | / | 0.02 / 0.35 | / | / | 5.88E-10 | 8.07E-08 |
| *p*-value 7 | / | / | 0.02 | 0.59 | 4.64E-09 | 4.58E-10 |
| *p*-value 8 | / | / | 0.01 | 0.24 | 9.68E-08 | 2.29E-08 |
| *p*-value 9 | / | / | 0.02 | 0.35 | 1.86E-08 | 2.97E-09 |
| growth rate (cm/d) ± SD | 0.68 ±0,03 | 0.63 ±0,04 | 0.70 ±0,04 | 0.66 ±0,01 | 0.66 ±0,02 | 0.64 ±0,02 |
| growth distance (cm)± SD | 15.4 ±1.7 | 30.6 ±10.2 | 24.3 ±9.1 | 30.8 ±7.7 | 57.9 ±20.5 | 67.6 ±26.6 |
| biological replicates | 25 | 34 | 26 | 15 | 22 | 21 |
